# A Novel Phosphatase Reverses the Leloir Pathway to Promote Tagatose Synthesis from Glucose

**DOI:** 10.1101/2025.07.29.665981

**Authors:** Aaron M. Love, Christopher G. Toomey, Abhishek Kumar, Sukesh Narayan Kashyap, Dhinesh Kumar Santhamoorthy, Likith Muthuraj, Hannah L. Lynch, Parayil Kumaran Ajikumar, Pravin Kumar R., Nikhil U. Nair, Christine N. S. Santos

## Abstract

D-Tagatose is a natural, low-calorie, rare monosaccharide that has gained substantial interest as an alternative sweetener. It offers numerous benefits, including a low glycemic index, prebiotic properties, and the ability to lower blood sugar levels. Currently, tagatose is primarily produced industrially through the isomerization of galactose, utilizing both chemical and enzymatic catalysis. While these established processes can produce tagatose, they are inefficient and expensive. Recent works have demonstrated alternative biosynthetic routes to produce D-tagatose but suffer from low theoretical yield and/or reliance on expensive feedstocks. This study demonstrates a novel biochemical pathway to produce D-tagatose directly from glucose using a whole-cell process with *Escherichia coli*. This process is distinct from other biosynthetic schemes and utilizes a newly discovered galactose-1-phosphate-selective phosphatase to drive the native Leloir pathway in reverse and synthesize the substrate D-galactose directly from D-glucose. Our analysis of the phosphatase reveals how an ensemble of intermolecular hydrogen bonds governs substrate specificity. By co-expressing this phosphatase and an L-arabinose isomerase in a modified strain background, we demonstrate production of tagatose directly from glucose. In initial studies, we generated ∼10.5 g/L galactose from 30 g/L glucose (35 % yield) while also producing > 1 g/L tagatose. This demonstrates the feasibility of a novel approach to tagatose production in vivo with a theoretical pathway yield of 94.9 %, which is substantially higher than previously proposed tagatose biosynthetic schemes.

## INTRODUCTION

In recent years, there has been growing interest in sourcing alternative natural sweeteners to replace both traditional sugars with high glycemic indices, as well as artificial sweeteners that have been under increasing scrutiny. D-tagatose (tagatose) is one such rare natural sweetener that is 92 % as sweet as sucrose when compared in 10 % solutions (Kim, 2004), and contains approximately one-third of the calories (Levin, 2002). Given its similar physical properties to sugar and GRAS approval by the FDA, tagatose is an attractive sweetener candidate to use in a variety of consumer products (Vera and Illanes, 2016). As with many low-calorie natural sweeteners, large-scale production of tagatose is limited by its scarcity in nature, as well as inefficient synthesis methods that suffer from low yields and purity, or feedstock limitations (Oh, 2007).

Tagatose is a C4 epimer of fructose and the ketose isomer of galactose, making it chemically similar to common commodity sugars and accessible through a minimal number of chemical transformations. It was initially synthesized economically by isomerization of galactose derived from inexpensive whey powder using calcium catalysts, in a process patented in 1992 (Kim, 2004). This route, however, suffers several disadvantages, including complex purification steps and the formation of chemical waste byproducts that hinder its implementation (Oh, 2007). Accordingly, there have been several proposed routes to make tagatose through biological production methods. The three most promising methods published include i) direct isomerization of galactose to tagatose with L-arabinose isomerase (LAI) (Bober and Nair, 2019; Cheetham and Wootton, 1993; Rai et al., 2021), ii) redox reactions through a galactitol intermediate (Liu et al., 2019; W. Liu et al., 2023), or epimerization of fructose to tagatose with (Lee et al., 2017b) or without phosphorylation-dephosphorylation steps (Shin et al., 2020).

While existing biosynthetic methods have been promising, they all suffer from fundamental limitations. The use of LAI as either a recombinant enzyme process or whole cell bioconversion has been engineered and improved (Bober and Nair, 2019), but the approach relies on a galactose feedstock, which is conventionally derived from the disaccharide lactose, consisting of glucose and galactose. Thus, there is a 50 % loss in convertible substrate that immediately limits the potential yield of any pathway relying on lactose, which includes the reported pathway using galactitol (Liu et al., 2019). Given that lactose has historically been more expensive than alternatives such as glucose (Zumbé et al., 2001), or at best has similar price ranges (USDA Economic Research Service, 2024; USDA Economics Statistics and Market Information System, 2024), using only 50 % of the substrate product formation is far from ideal (Fan et al., 2025). Fructose can be epimerized directly to tagatose, but it suffers from similar problems with poor yield at equilibrium (Shin et al., 2020). Conversely, the phosphorylation-dephosphorylation strategy enables epimerization of fructose-6-phosphate to tagatose-6-phosphate with yields over 90% by driving the epimerase reaction equilibrium towards the desired product (Lee et al., 2017b), but it does not work efficiently in vivo due to inhibition of the epimerase by fructose-1,6-bisphosphate (Lee et al., 2017a), which ultimately limits economic feasibility, given that in vitro processes are substantially more expensive than using whole cells (Claassens et al., 2019).

A more desirable theoretical approach for tagatose biosynthesis would use whole cells without the need for expensive enzyme purification, combined with a biosynthetic pathway enabling production of tagatose from inexpensive feedstocks like glucose while utilizing all the available substrate for production. In this work, we demonstrate this approach by engineering a strain of *E. coli* that produces high levels of galactose directly from glucose through reversal of the Leloir pathway. Borrowing from principles exemplified by the phosphorylation-dephosphorylation approaches previously used to drive high yield of rare sugar biosynthesis, we identify a novel galactose-1-phosphatase (Gal1Pase) capable of shifting the natural equilibrium of the Leloir pathway. We perform rigorous computational analysis of Gal1Pase using a combination of molecular dynamics (MD), quantum mechanics/molecular mechanics (QM/MM), and well-tempered metadynamics simulations to explain how galactose is discriminated so well from its structurally similar, and more abundant, C4 epimer glucose. Without substantial optimization, we build strains capable of converting 35% of fed glucose to galactose, producing over 1 g/L tagatose in whole cell cultures of *E. coli*, demonstrating an attractive new route to this rare sugar.

## RESULTS

### A screen for Gal1P-specific phosphatases identifies a novel enzyme from slime mold

The simplest theoretical tagatose production pathway using glucose as a substrate is through C4 hydroxyl epimerization and aldose-ketose isomerization (glucose ➔ galactose or fructose ➔ tagatose). This route, however, is kinetically and thermodynamically unfavorable, and the use of phosphorylation-dephosphorylation steps can circumvent these issues. We initially explored this approach, where tagatose-6-phosphate (Tag6P) can be dephosphorylated to drive the reaction to completion (S. Liu et al., 2023). (**Fig. S1**). Since UDP-glucose-4-epimerases can act on free glucose (Kim et al., 2011), we thought they may also be capable of converting glucose-1-phosphate (G6P) to galactose-1-phosphate (Gal1P) – given the higher structural similarity between the nucleoside and phosphosugar compared to the free sugar. A subsequent enzyme would then be required to isomerize Gal1P to Tag1P, which has, unfortunately, not been documented. Instead, isomerization occurs naturally between galactose-6-phosphate (Gal6P) and Tag6P (Kim, 2004). Since Gal1P, not Gal6P, is an intermediate in our proposed pathway, it would require a phosphomutase step to isomerize Gal1P to Gal6P. This reaction has been reported in the literature but is strongly inhibited by G1P (Mukherjee et al., 2018). Initially, we investigated a panel of diverse epimerases with substantial phylogenetic variation to identify candidates with G1P to Gal1P activity (**Fig. S2**). Many of these were selected from prior reports of epimerases that catalyze similar reactions (Nam et al., 2019). While we observed high activity from most epimerases for the conversion of UDP-galactose to UDP-glucose, we did not observe any detectable activity for the interconversion between Gal1P and G1P (**Fig. S3** and **S4**).

Given this disappointing result, we decided to instead leverage *E. coli* GalET to generate Gal1P. While the native activity of this enzyme pair favors formation of G1P during galactose catabolism (in the Leloir pathway), it is reversible and can be driven towards galactose production by selective dephosphorylation of Gal1P (**Fig. 1A–C**). In this case, glucose could be directly converted to galactose, which could then be transformed to tagatose using the well-characterized promiscuous LAI enzymes (Baptista et al., 2021; Bober and Nair, 2019; Kim, 2004; Shin et al., 2022). This strategy was predicted by pathway analysis to be even more effective with the combined genetic knockouts of Δ*pgi* and Δ*galK* to divert metabolic flux away from glycolysis and prevent the re-phosphorylation of free galactose, respectively (**Fig. 1A**). When combined, the reactions reverse the Leloir pathway and are expected to be thermodynamically favorable for the formation of intracellular galactose and tagatose (**Fig. 1B and C**). This pathway also has a theoretical yield of 94.9 %, only requiring minimal expenditure of glucose substrate for ATP production (**Fig. S5**).

**Fig. 1.**
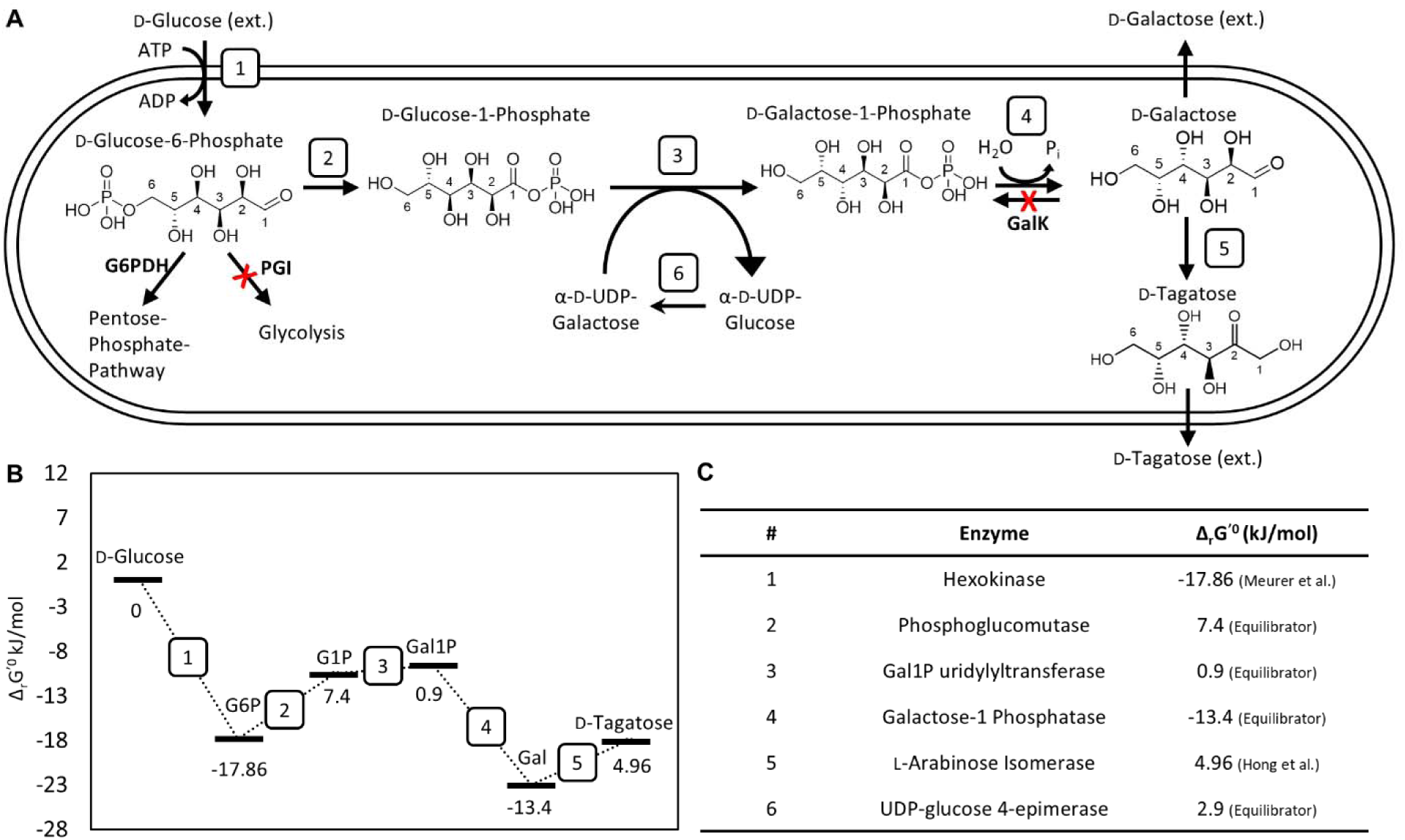
Proposed biosynthetic route from glucose to tagatose by reversing the Leloir pathway. **(A)** Implementation of this pathway involves 6 reactions and inactivation of 2 native genes. (**B**) Approximate standard Gibbs free energy change for each reaction step numbered in **A** between glucose and tagatose using the reverse Leloir pathway. Y-axis values represent the net change in free energy due to each reaction starting from glucose as a basis. (**C**) Table of enzymes catalyzing each step represented in **A**, and the corresponding free energy change in kJ/mol for the reaction direction shown in **A**. Δ*_r_*G’° represents the change in Gibbs free energy due to the indicated reaction under standard conditions. The values for hexokinase and LAI were taken directly from the literature (Meurer et al., 2017) and (Hong et al., 2007) respectively. Others were calculated using the eQuilibrator tool (http://equilibrator.weizmann.ac.il) (Beber et al., 2022).

All the enzymes required for the biosynthetic scheme in **Fig. 1A** had been characterized except for the Gal1P phosphatase (Gal1Pase). While there had been instances of promiscuous phosphatases acting on Gal1P (Parthasarathy et al., 1997) or specific L-galactose-1-phosphate (Laing et al., 2004), there are no documented enzymes with a strong preference for Gal1P over G6P. Accordingly, we screened several phylogenetically diverse phosphatases to identify candidates with the desired activity and specificity towards Gal1P (**Fig. 2A–B**). The average sequence identity between any 2 phosphatases screened was only 23 % (**Fig. S6**). In total, we selected 9 candidates, codon optimized them and expressed them in *E. coli* BL21(DE3). All phosphatases expressed well based on SDS-PAGE analysis (**Fig. 2C**), except for the one from *B. subtilis*, which was thereafter excluded from further analysis.

**Fig. 2.**
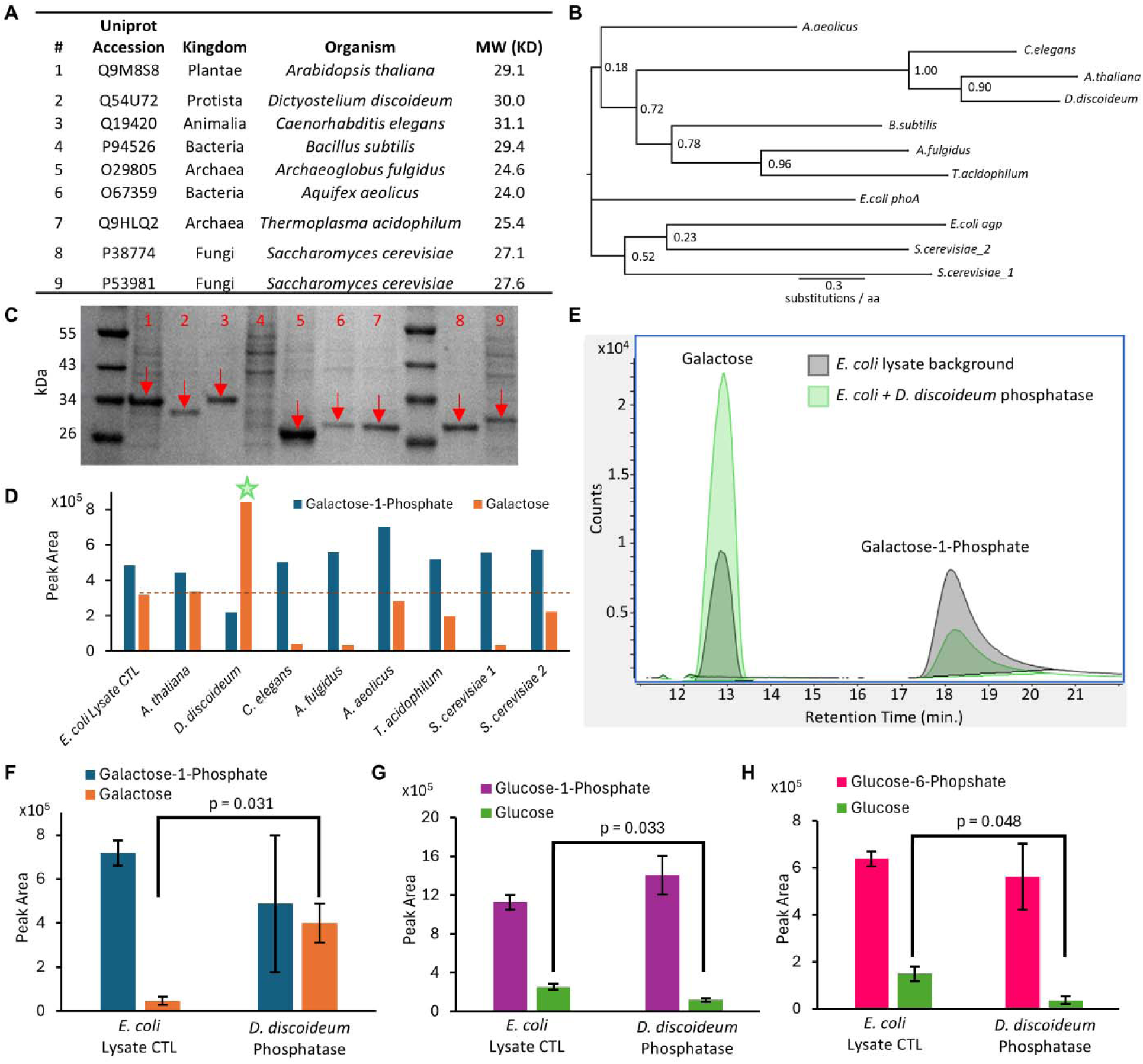
Discovery and characterization of Gal1P-specific phosphatase from *D. discoideum* (DdGal1Pase). (**A)** Table summarizing phosphatases selected for screening. (**B)** Phylogenetic tree of phosphatases screened, highlighting their evolutionary distance. *E. coli* phosphatases included for reference. Support values are indicated at branch points. (**C)** SDS-PAGE results from lysates obtained after phosphatase expression in *E. coli* BL21(DE3) loaded with normalized total protein. Lane numbers also correspond to the enzyme source indicated in (**A**). Arrows indicate recombinant enzymes. Only the *B. subtilis-*derived enzyme (lane 4) did not show clear soluble expression. (**D**) Evaluation of enzyme assay with Gal1P substrate. Shown are quantified raw peak areas from chromatograms for Gal1P and galactose after incubation with different phosphatase-containing lysates. Star (★) indicates activity higher than control. (**E**) LC-MS chromatogram showing Gal1Pase activity – depletion of Gal1P and evolution of galactose – by the enzyme derived from *D. discoideum*. Substrate specificity screen comparing phosphatase activity of lysates containing DdGal1Pase on (**F**) Gal1P, (**G**) G1P, or (**H**) G6P. Only Gal1P shows higher activity with DdGal1Pase than with the lysate control. Bars represent mean ± SD, all p values represent two-tailed t-tests, *n = 2*.

We first assessed phosphatase activity in crude lysates by feeding Gal1P and looking for the evolution of galactose. Of all the phosphatases we screened, only the variant from the slime mold *Dictyostelium discoideum* (DdGal1Pase) exhibited activity towards Gal1P higher than the WT *E. coli* lysate control (**Fig. 2C–E**). Since reactions were normalized to total protein, several lysates exhibit lower activity than the control, likely due to a reduction in abundance of native phosphatases upon overexpression of recombinant enzymes. We re-screened DdGal1Pase relative to the *E. coli* lysate control for substrate specificity using Gal1P, G1P, and G6P.

DdGal1Pase exhibited significant activity on Gal1P (p = 0.031) but did not show activity over background on either G1P or G6P (**Fig. 2F–H, Fig. S7**). This result is remarkable given the similarity between glucose and galactose, only differing by the C4 hydroxyl stereochemistry, marking the first description of a phosphatase with this narrow substrate range.

### DdGal1Pase substrate specificity drives reversal of the Leloir pathway

Following the discovery of a Gal1P-specific phosphatase, we assessed its performance in aiding growth-coupled conversion of galactose to tagatose. We constructed an initial strain with *galK* and *galM* deleted to promote the accumulation of galactose and conversion to tagatose (Kim et al., 2008) and prevent futile re-phosphorylation. After transforming this strain with a plasmid encoding DdGal1Pase and a previously characterized L-arabinose isomerase (LAI) from *Bacillus coagulans* (BcLAI) (Mei et al., 2016), we cultured strains containing 30 g/L galactose and quantified the evolution of glucose and tagatose. The unmodified *E. coli* control metabolized substantial quantities of the fed galactose and accumulated g/L quantities of glucose, presumably through the Leloir pathway. Consistent with previous reports, the Δ*galKM* strain used galactose only to produce tagatose at higher levels than the WT *E. coli* control (**Fig. 3A**). In this case, we observed 1.8 g/L tagatose compared to only 0.4 g/L in the WT strain. Interestingly, small quantities of tagatose formed without BcLAI complementation, which may be attributed to leaky expression of the native LAI (*araA*). Overall, tagatose production with this LAI was lower than in previous reports (Mei et al., 2016) likely due to differing conditions (e.g., growth temperature).

**Fig. 3.**
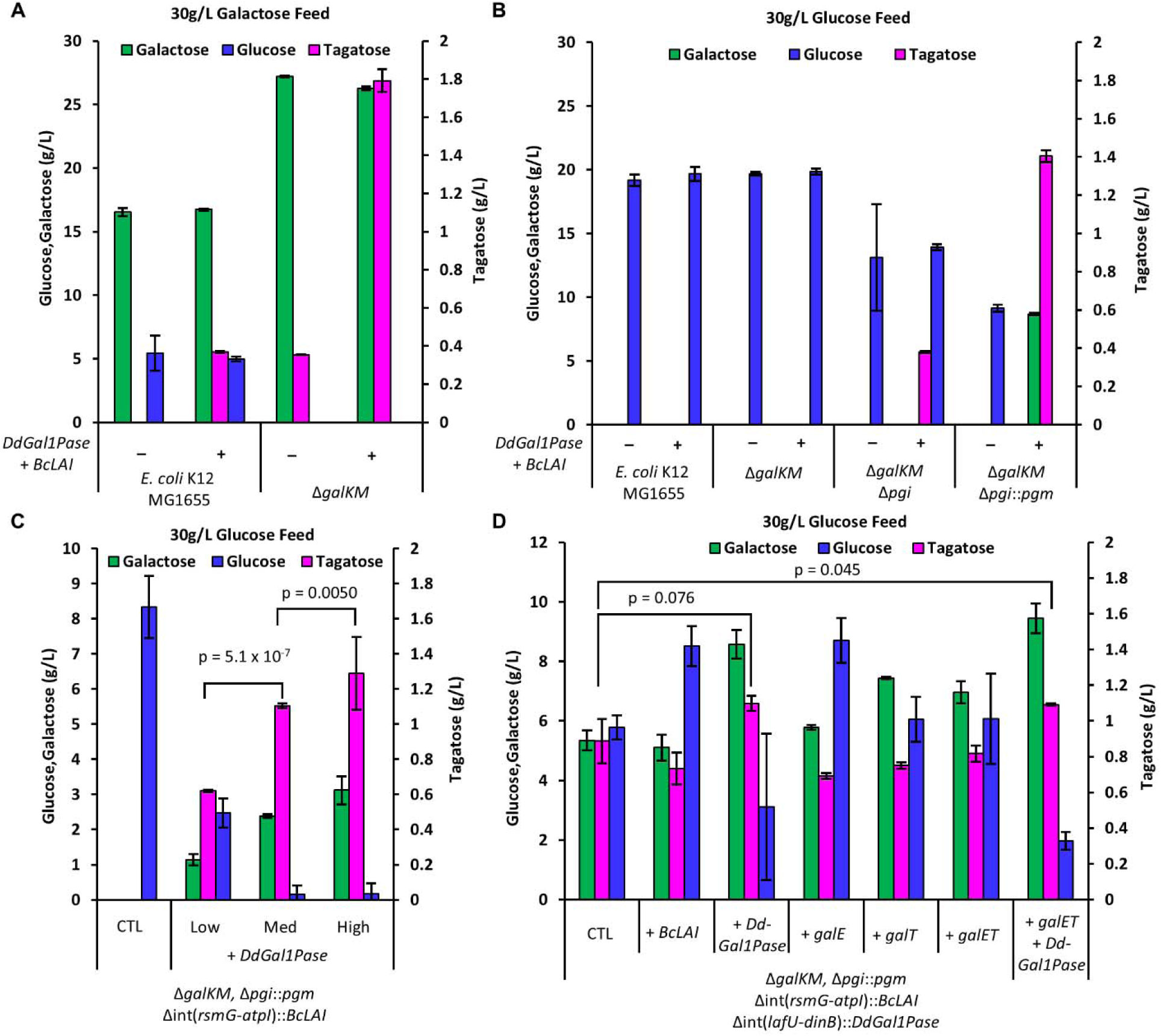
**Whole cell biosynthesis of tagatose**. Expression of *DdGal1Pase* and *BcLAI* from plasmid (indicated with + or –) in 4 different *E. coli* strain backgrounds fed (**A**) 30 g/L galactose or (**B**) 30 g/L glucose. The Δ*galKM* modification improves conversion of galactose to tagatose and reduces galactose consumption by native metabolism. Reversal of the Leloir pathway is enabled by *pgi* deletion and PGM overexpression combined with the Gal1P-specific DdGal1Pase. (**C**) The Leloir pathway reversal is dependent on DdGal1Pase expression level modulated by changing RBS strength, highlighting its key role in the pathway. (**D**) Complementation to assess the limitation of various genes in a fully integrated tagatose production strain with a *pgi* deletion and chromosomal overexpression of *pgm, BcLAI*, and *DdGal1Pase*. Only DdGal1Pase overexpression significantly improves mean tagatose production. Bars represent means ± SD, *n = 3*, all p values represent two-tailed t-tests.

Next, we assessed whether strains co-expressing DdGal1Pase and BcLAI could be used to convert glucose directly to galactose and tagatose. When grown in 30 g/L glucose, both the WT and the Δ*galKM* (control) strains metabolized glucose without producing any detectable galactose or tagatose (**Fig. 3B**). To push glucose into the reverse Leloir pathway, we deleted *pgi* to block glycolysis. This improved glucose conversion and yielded low levels of tagatose (∼0.4 g/L). Finally, overexpression of *pgm* was key to driving high levels of galactose and tagatose formation. Remarkably, the Δ*galKM* Δ*pgi*::*pgm* strain formed 8.7 g/L galactose and 1.4 g/L tagatose without any other optimization.

We then investigated the role of DdGal1Pase activity in driving tagatose production. After chromosomally integrating *BcLAI* into the Δ*galKM* Δ*pgi*::*pgm* strain, we complemented *DdGal1Pase* on the pBAC plasmid with three ribosome binding sites (RBSs) of increasing strengths. **Fig. 3C** shows a clear dose-dependent response to increasing levels of phosphatase expression on tagatose and galactose production. This result demonstrates that galactose production from glucose is strongly dependent on DdGal1Pase activity. To evaluate limiting components of the pathway, we also chromosomally integrated *DdGal1Pase* into the Δ*galKM* Δ*pgi*::*pgm BcLAI* strain. We then supplemented each gene in the pathway episomally, one at a time (**Fig. 3D**). Results indicate that DdGal1Pase is the rate-limiting enzyme in the pathway, as complementing it increases galactose titers from 5.4 to 8.6 g/L, and tagatose from 0.9 to 1.1 g/L. Additional BcLAI expression did not improve tagatose production, indicating the enzyme kinetics are probably not limiting. It also appears that the enzymes GalE and GalT are not rate-limiting for galactose production, though there is some benefit to additional *galT* overexpression. Despite the expected carbon catabolite repression on the Leloir pathway when growing on glucose, it appears the *galET* genes are already expressed in sufficient quantity from the modified strain background, which could be due to catabolite repression relaxation that can occur in to Δ*pgi* strains (Fox and Prather, 2020; Shiue et al., 2015). Since increased galactose production by increasing DdGal1Pase overexpression leads to more tagatose, galactose secretion probably limits tagatose production. Further, the low reaction temperature is unfavorable for galactose to tagatose isomerization based on previous reports (Bober and Nair, 2019).

We attempted to address thermodynamic equilibrium constraints in the system by modulating the reaction temperature or the sugar partitioning across the cellular membrane. Since higher temperatures favor the isomerization reaction towards tagatose (Bober and Nair, 2019; Rai et al., 2021; Xu et al., 2018), we evaluated performance from our fully integrated tagatose pathway strain (from **Fig. 3D**). with or without a temperature shift to 60 °C for 24 h (**Fig. 4A–B**). The temperature-shifted cells exhibited substantially higher tagatose formation, but little additional glucose consumption compared to controls. This indicates that while additional tagatose was produced from existing galactose via LAI activity, glucose metabolism ceased after cells were subjected to lethal temperatures. Mn^2+^ supplementation to promote LAI activity (Mei et al., 2016) did not improve tagatose production (**Fig. S8**). Selective partitioning of galactose and tagatose across the cell membrane has been shown to improve equilibrium conversion (Bober and Nair, 2019; Kim et al., 2008). *E. coli* has several known and putative sugar transporters that could help to differentially partition sugars to favor tagatose formation (**Table S1**). We targeted 12 genes for deletion to study their effect on tagatose formation in the fully integrated tagatose pathway strain (Carreón-Rodríguez et al., 2023; El Qaidi et al., 2009; Kim et al., 2008; Koita and Rao, 2012; Tong et al., 2020). Of these, the deletion of *ydeA* exhibited a 1.66-fold increase in tagatose production, while the deletion of the exporter *setC* had a lower but significant 1.18-fold improvement (**Fig. 4C**). Overexpression of *ydeA* decreased tagatose production, as expected. It also unexpectedly seemed to interfere with glucose uptake (**Fig. S9**). Unfortunately, the combination of Δ*ydeA* and Δ*setC* did not further improve tagatose production (**Fig. S10**). We further characterized the Δ*ydeA* strain by testing it with 30 g/L or 5 g/L glucose feeding, as well as a temperature shift to 50 °C. In this case, 50 °C was used to improve equilibrium conversion to tagatose while preserving the activity of BcLAI for a longer duration. We again see improved conversion of galactose to tagatose in the Δ*ydeA* context and a positive impact of shifting to 50 °C after 24 h. of growth at 37 °C. Interestingly, the Δ*ydeA* strain is not differentiated with lower 5 g/L glucose feeding. This could be due to lower metabolic flux towards galactose, thus lowering intracellular galactose concentrations and reducing the relative efflux of galactose vs. conversion to tagatose. In this hypothetical case, the impact of deleting a galactose efflux pump would be reduced. This result suggests that a combination of optimal glucose feeding combined with transporter engineering could further increase tagatose yield.

**Fig. 4.**
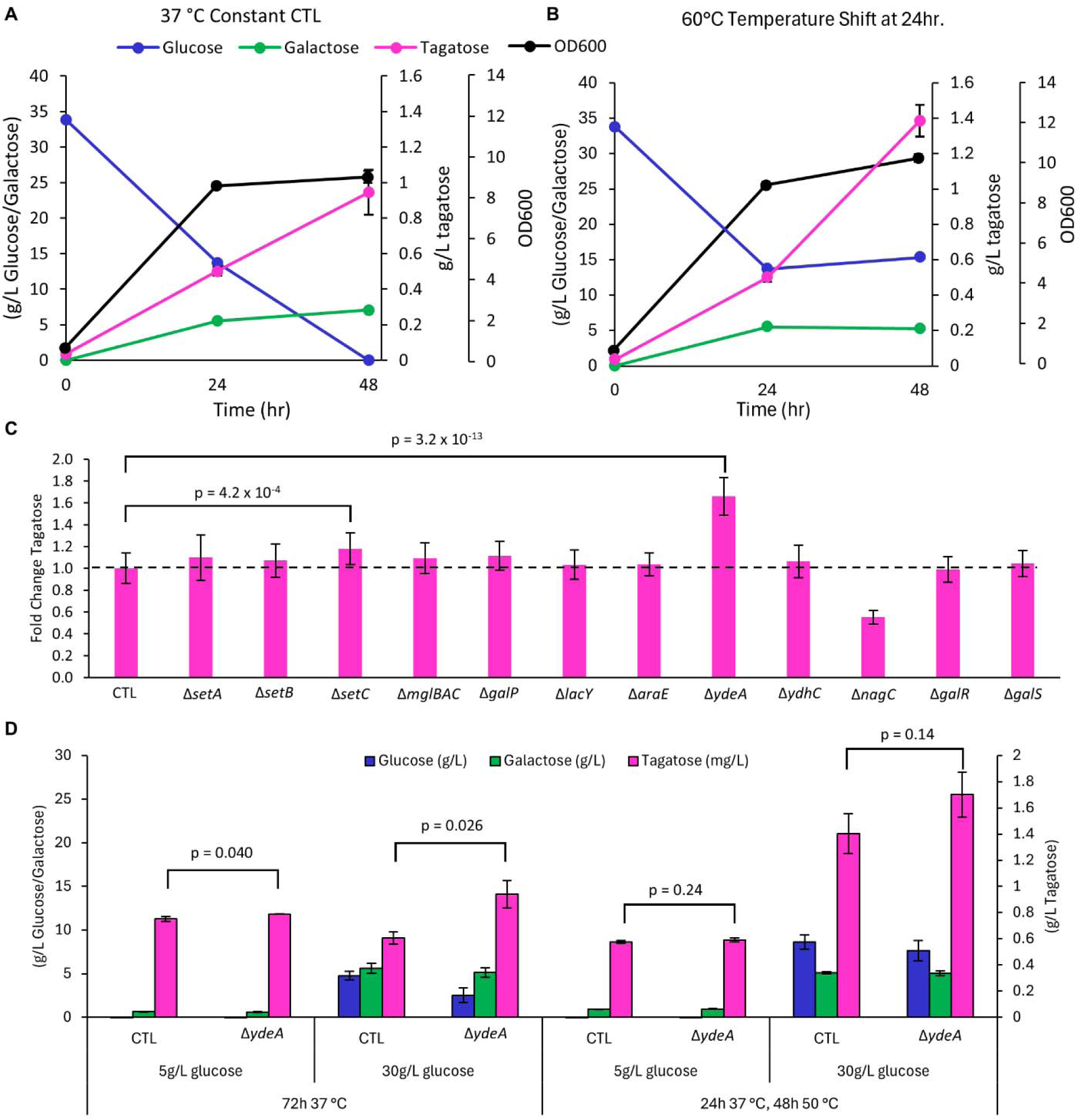
Thermodynamic controls improve tagatose production. Temperature change to (**A**) 37 °C or (**B**) 60 °C after 24 h reveals that higher temperature favors tagatose production. (**C**) A transporter deletion screen reveals that deletion of the putative sugar exporter, *ydeA,* improves tagatose production. (**D**) Characterization of Δ*ydeA* mutant strain with either 5 g/L or 30 g/L glucose feeding with or without a shift from 37 °C to 50 °C after 24 h. Bars represent means ± SD, n = 3, all p values represent two-tailed t-tests.

Given how critical terminal phosphatase substrate specificity is to this pathway, we investigated how the structure of DdGal1Pase relates to its stringent discrimination of different substrates. To explore the unique specificity of DdGal1Pase on Gal1P, we tested homologs that were similar in sequence. A phylogenetic analysis revealed 5 relatively closely related sequences that were all > 70 % sequence identity to DdGal1Pase compared to the next closest homolog that was only < 50% identical. Given this distinctly similar set of 5 sequences (**Fig. S11A**), we re-coded them for expression in the Δ*galKM* Δ*pgi*::*pgm BcLAI* background. We also overexpressed two native sugar phosphatases, *agp* and *phoA,* from *E. coli* to verify that the observed Gal1P specificity is essential for Leloir pathway reversal. Our data confirms that DdGal1Pase drives Leloir pathway reversal compared to Agp and PhoA, and 4 of the 5 tested homologs also exhibit similar Gal1P phosphatase activity in vivo (**Fig. S11B**).

### Molecular simulations indicate subtle interactions control DdGal1Pase substrate specificity

To further characterize the unique specificity for gal1P exhibited in this sequence space, we modeled DdGal1Pase with G1P and Gal1P in the active site. DdGal1Pase was structurally modeled using AlphaFold (Abramson et al., 2024), as DdGal1Pase showed a maximum sequence identity of 41.1% with reposited crystallographic structures (PDB ID: 2QFL, inositol-1-monophosphatase from *E. coli*), and the modelled structure of DdGal1Pase showed a predicted template modelling score (pTM) of 0.93 and a predicted local distance difference test (pLDDT) of > 90 in every region, except at the termini and residues 30–46 (**Fig. S12A**). The stereochemical assessment of DdGal1Pase structure showed a Ramachandran favored region of 97.41% (**Fig. S12B**). The sequence alignment of DdGal1Pase with the known protein sequence database revealed it as an inositol monophosphatase enzyme with three identical motifs (A, B, and C), similar to the human inositol monophosphatase (Bone et al., 1994), and two catalytic and one non-catalytic Mg^2+^ ions in the active site (**Fig. S13**) (Bone et al., 1994; Wang and Hirao, 2013). The derived DdGal1Pase substrate complexes (with either Gal1P or G1P) were selected based on the hydrogen bonding of substrate hydroxyl groups to active site residues (D93, G94, T95, A196, E213, and D220), as previously reported for the inositol moiety in inositol monophosphates (Atack et al., 1995; Bone et al., 1994) (**Fig. 5A–B**). We observed an obvious difference in the orientation of E213 and hydrogen bonding distance with the C4 epimeric substrates, Gal1P and G1P, suggesting a preference for Gal1P (**Fig. 5A–B**). Our analysis also revealed that the catalysis-initiating residue E71 doesn’t interact directly with the substrate molecule but initiates the reaction through a water molecule interacting with _1_Mg^2+^ ion (**Fig. 5C**).

**Fig. 5.**
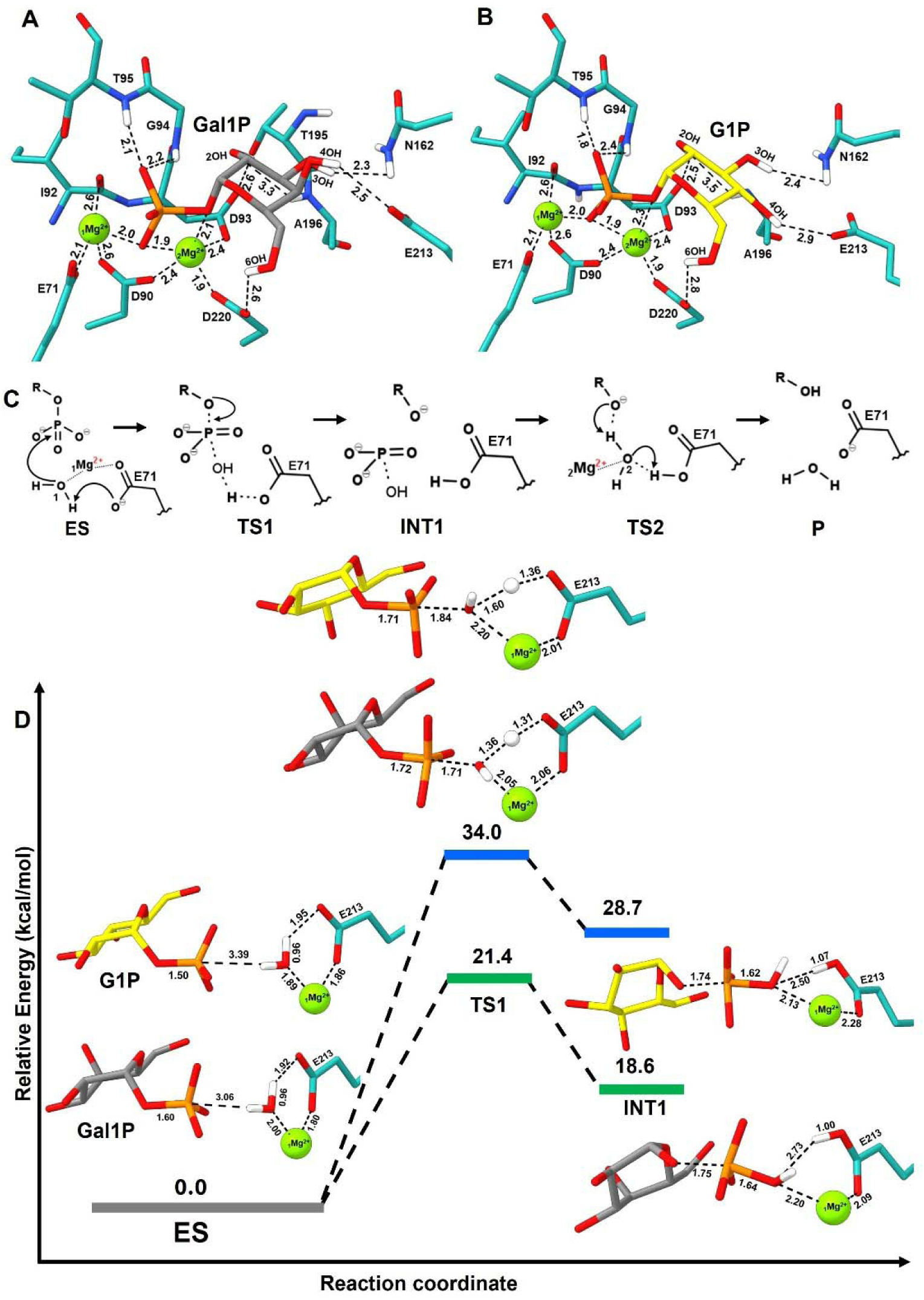
MD and QM/MM analysis of DdGal1Pase. Gal1P demonstrates closer interactions with active site residues and Mg²D ions compared to G1P. The derived enzyme substrate (ES) complex of DdGal1Pase with substrates (**A**) Gal1P (grey) and (**B**) G1P (yellow). (**C**) The plausible reaction mechanism of DdGal1Pase, highlighting the essential role of water molecules in the phosphate hydrolysis reaction mechanism with two Mg^2+^ ions. The _1_Mg^2+^ and _2_Mg^2+^ ions and water molecules are numbered according to their involvement in catalysis. (**D**) The calculated energy profile for the phosphate hydrolysis of substrates Gal1P and G1P with their optimized structures of transition states (TS) and intermediates (INT1) along the reaction pathway. Non-polar hydrogen atoms have been omitted for clarity. The Gal1P (grey) and G1P (yellow) substrates are shown as ball and stick models, the Mg^2+^ ions are shown as green spheres, and key active site residues are shown in light sea green. The atomic interactions are shown with black dotted lines with distances in Å. G1P = glucose-1-phosphate, Gal1P = galactose-1-phosphate.

We then conducted molecular dynamics (MD) simulations to investigate the role of the active site residues of DdGal1Pase that control the stringent selectivity for Gal1P over G1P. Root mean square deviation (RMSD) analysis indicates that the *apo* form of DdGal1Pase exhibits the greatest structural deviation, signifying increased conformational flexibility and dynamic instability in the absence of substrate-induced stabilization. Upon binding with Gal1P, the preferred substrate, Ddgal1Pase demonstrates the lowest RMSD, suggesting a more stable and compact conformation likely associated with a catalytically competent state. G1P binding results in intermediate RMSD values, reflecting partial stabilization of the enzyme structure, possibly due to suboptimal interactions within the active site (**Fig. S14**). A conserved characteristic among IMPase family enzymes, including archaeal IMPase and DdGal1Pase, is the presence of mobile loops proximal to the active site (Li et al., 2010). These mobile loops frequently exhibit substrate-induced dynamic behavior to facilitate substrate accommodation, thereby achieving the enzyme’s catalytic state. Indeed, our modelling suggests these regions could be implicated in discriminating between Gal1P over G1P (**Fig. S15A**). Additionally, these loops are integral to coordinating the third metal ion, stabilizing the nucleophilic water molecule, and optimizing substrate positioning. Their flexibility and acidic nature are crucial for sustaining catalytic efficiency (**Fig. S15B–C**). These findings underscore the substrate-specific stabilization effect of Gal1P and suggest a molecular basis for the enzyme’s substrate selectivity in DdGal1Pase.

We also found notable differences in non-covalent interactions through the analysis of hydrogen bonding distances between 2OH, 3OH, 4OH, and 6OH on the substrate pyranose rings and D93, N162, E213, and D220, respectively (**Fig. S16**). These hydrogen bonding interactions are essential for stabilizing substrate binding and contribute to the stringent selectivity of DdGal1Pase for Gal1P. An assessment of hydrogen bond distances revealed substrate-dependent variations across multiple residues. Namely, the hydrogen bonding interaction between Gal1P and D220 at the 6OH position demonstrated a favorable distance conducive to hydrogen bond formation. In contrast, G1P exhibited a greater distance, which is unfavorable for hydrogen bonding. With D93 and 2OH, Gal1P displayed an unfavorable distance for hydrogen bonding, whereas G1P showed a favorable distance. The A196–2OH hydrogen bond length did not show any significant difference between the two substrates. Importantly, E213–4OH and N162–3OH hydrogen bonding interactions occurring at the epimer and epimer-preceding atoms, respectively, showed shorter hydrogen bond distances in Gal1P as compared to the G1P (**Fig. S17**). Collectively, these observations suggest that the hydrogen bonding pattern differs significantly between Gal1P and G1P, influencing their interactions with the enzyme’s active site. Importantly, the hydrogen bonding residues D220, N162, and E213 contribute to preferential binding for catalytic activity on Gal1P. Conversely, for G1P, the most favorable hydrogen bonding distance was D93–2OH, which favors the non-catalytic positioning of G1P in the active site, and leads to the enzyme’s poor activity toward G1P.

To further probe DdGal1Pase specificity, we conducted a QM/MM study to investigate the energetic feasibility of the reaction involving either Gal1P or G1P. The QM/MM mechanism of DdGal1Pase for Gal1P and G1P was elucidated based on the known catalytic mechanism of inositol monophosphatase with two metal ions (Wang and Hirao, 2013). The DdGal1Pase was modelled with three water molecules coordinated by Mg^2+^ ions. The proposed two-metal-ion mechanism is initiated by the catalytic residue E71 acting as a base. In this mechanism, a water molecule bound to _1_Mg² donates its proton to E71, generating a hydroxide ion. The formed hydroxy acts as a strong nucleophile that attacks the phosphate of the substrates resulting in hydrolysis (**Fig. 5C**). The reaction mechanism of Gal1P and G1P showed a clear distinction in the energies of transition state 1 (TS1) and intermediate state 1 (INT1) between Gal1P and G1P (**Fig. 5D**). The QM/MM mechanism of DdGal1Pase has been elucidated up to the phosphate hydrolysis step (INT1). Subsequent steps are not the contributing factors for substrate specificity in DdGal1Pase. In TS1, the proton transfer from water to E71, coupled with the nucleophilic attack of the resulting hydroxyl on the phosphate group, showed an energy of 21.4 kcal·mol^−1^ for Gal1P, while G1P showed a significantly higher energy of 34.0 kcal·mol^−1^. Further, the protonated E71 and hydroxylated phosphate group in INT1 showed an energy of 18.6 kcal·mol^−1^ for Gal1P and 28.7 kcal·mol^−1^ for G1P (**Fig. 5D**). QM/MM simulations reveal that the Gal1P-bound enzyme complex achieves a faster and more consistent reduction in the distance between the phosphate and the water nucleophile compared to the G1P-bound system (**Fig. S18A**). Specifically, Gal1P reaches a reactive conformation in fewer reaction coordinates, indicating a more favorable and native-like binding pose for catalysis by DdGal1Pase. In contrast, G1P requires nearly three times the optimization steps to achieve a similar conformation, suggesting less efficient activation. Supporting this notion, the _1_Mg² –O14–P angle stabilizes more quickly for a geometrical alignment required for nucleophilic attack by hydroxyl ion to phosphate (**Fig. S18B**). This angle remains consistent post-reaction in the Gal1P system, further validating that Gal1P forms a catalytically competent geometry more readily than G1P.

The hydrogen bonding analysis and QM/MM analysis highlighted the selectivity of Gal1P and G1P. Additionally, we performed a comprehensive evaluation to assess the catalytic distance (coordinate bonds) stability of the Mg^2+^ ions and the diffusion of water molecules to these ions, which are integral to the catalytic process. The _1_Mg² ion adopts a stable six-coordinate bond geometry with residues D90, E71, and the backbone of I92, as well as the substrate’s phosphate oxygen atoms. Similarly, the _2_Mg² ion forms a six-coordinate bond geometry with residues D90, D93, D220, and the substrate’s phosphate oxygen atoms (**Fig. S19**). A subtle, substrate-dependent shift in the I92 backbone coordination is observed between the Gal1P- and G1P-bound states with _1_Mg² ion (**Fig. S19A**) and with _2_Mg² ion the coordinate bond of O6 atom of Gal1P exhibits a greater distance compared to the G1P complex forms (**Fig. S19B**). In the *apo* form, the non-catalytic _3_Mg² ion forms a stable coordinate bond (<2.0 Å) with the catalytic residue E71 (**Fig. S19C**). This interaction prevents the dissociation of the _3_Mg² ion from the active site and contributes to maintaining the structural integrity of the active site for the substrate accommodation (Lu et al., 2012) and the catalytic distance stability of _1_Mg^2+^ and _2_Mg^2+^ ions remains unaffected by the presence of substrates (Gal1P and G1P) within the active site. The presence and dynamic behavior of water molecules in proximity to Mg² ions play a critical role in facilitating phosphate hydrolysis. Specifically, coordination of water molecules with _1_Mg² ion promotes the hydrolysis reaction, while a _2_Mg² ion contributes to the protonation of the pyranose moiety post-hydrolysis. Moreover, these water molecules assist in satisfying the remaining coordination sites of the Mg² ions that are not occupied by active site residues (Wang and Hirao, 2013). To investigate the spatial distribution and likelihood of water molecule coordination around the Mg² ions, a radial distribution function (RDF) analysis was performed. The RDF provides a quantitative measure of the probability of finding catalytic water molecules within a radial distance characteristic of coordination bond formation (**Fig. S20**). The RDF analysis revealed that water molecules were consistently present within 2.0 Å of the _1_Mg² ion in both the *apo* and **DdGal1Pase**-Gal1P/G1P forms (**Fig. S20A**). In contrast, the _2_Mg² ion in the Gal1P-bound complex showed significantly reduced water presence within 2.0 Å, with no water molecules observed in the G1P-bound form indicating substrate-dependent water modulation and a more tightly coordinated active site around the _2_Mg² ion (**Fig. S20B**). In the *apo* form of DdGal1Pase, the _3_Mg² ion is coordinated by more than five water molecules within 2.0 Å, consistent with a six-coordinate bond geometry when including the interaction with residue E71 (**Fig. S20C**).

### Metadynamics reveals distinct energy minima favoring Gal1P specificity

In DdGal1Pase, the substrate is stabilized in the active site through a network of hydrogen bond interactions between the pyranose ring and surrounding residues. To better understand this network, we conducted an in-depth hydrogen bond occupancy analysis to probe the catalytic selectivity of DdGal1Pase towards Gal1P over G1P. The hydrogen bond occupancy analysis exhibited unique interactions of the active site residues with both substrates, Gal1P and G1P. Many of the active site residues including N162, G94, G194, T95, E213, K38, and G215 showed indistinguishable hydrogen bonding interactions with Gal1P and G1P, eliminating their role in substrate selectivity (**Fig. 6A**). A subset of active site residues exhibited transient hydrogen bond interactions with the substrates, defined here as cryptic interactions (mean hydrogen bond occupancy <1 %). While several residues formed hydrogen bonds to both substrates, others displayed substrate-specific interactions across the simulation trajectories. For instance, Y165, A196, W219, and D42 engaged in cryptic hydrogen bonding with both Gal1P and G1P. In contrast, K75 exhibited a unique interaction with Gal1P, whereas T96, G164, and H217 demonstrated hydrogen bonding exclusively with G1P (**Table S2**). These cryptic hydrogen bond interactions are unlikely to influence substrate discrimination significantly. However, a distinct pattern of hydrogen bond occupancy is evident between Gal1P and D220 (ANOVA, p = 0.030) and G1P and D93 (ANOVA, p = 0.034). Specifically, Gal1P exhibits a preferentially higher bond occupancy with D220 compared to G1P, whereas G1P demonstrates a preferentially higher bond occupancy with D93 compared to Gal1P (**Fig. 6A**). Structurally, D220 and D93 residues are spatially positioned in direct opposition to each other within the active site and are involved in the _2_Mg^2+^ ion binding. The two substrates are C4 epimers, differing in the orientation of the hydroxyl group, being axial in Gal1P and equatorial in G1P. Residue E213 stabilizes the C4 epimer, exhibiting approximately three times higher hydrogen bonding occupancy with Gal1P than with G1P (**Table S2**). E213 is positioned on a β-sheet that is connected via a loop to an α-helix containing D220 (**Fig. S21**). The Gal1P axial 4OH hydroxyl has a higher chance of hydrogen bonding with E213, which promotes favorable interactions between D220 and 6OH. In contrast, the equatorial 4OH of G1P has a lower chance of hydrogen bonding with E213, which promotes interaction between D93 and 2OH instead, but not with D220. This difference in interactions with D220 and D93 results in a 24.4° angular difference in the orientation of the pyranose rings of G1P and Gal1P (**Fig. 6B**). This result was further corroborated through the application of well-tempered metadynamics. This approach was employed to examine the conformational transitions in hydrogen bonding involving D220, D93, and the substrate 6OH and 2OH groups, respectively. Metadynamics simulations were performed using collective variables (CVs) defined as the distance between the center of mass (COM) between the two representative groups (D93–2OH and D220–6OH) (**Fig. S22**). These CVs were defined to understand the role of the two residues in substrate binding and subsequently determining the catalytic conformation. Results from the metadynamics simulations revealed a distinct energy minimum associated with each substrate. The Gal1P substrate, in its local energy minimum conformation, exhibited a collective variable CV distance of approximately 4 Å with both D93 and D220, indicating stable interactions with these residues (**Fig. 6C**). In contrast, the G1P substrate showed a broader distance distribution to D220, ranging from 5 to 6 Å with a distinct minimum centered at 5.5 Å, while maintaining a shorter and more consistent CV distance of less than 3.5 Å with D93 (**Fig. 6D**). This disparity suggests that G1P preferentially engages D93 over D220. These findings indicate that Gal1P adopts a more stable and specific induced-fit conformation within the active site of DdGal1Pase compared to G1P. To further explore the structural changes and interactions responsible for these differences, we extracted a representative structural complex from the minima to analyze the induced fit conformation through hydrogen bond interactions involving three hydroxyl groups of the substrates. In this induced fit conformation, the 2OH formed a hydrogen bond interaction with G94, the 4OH with E213, and the 6OH with D220, indicating a stable induced fit conformation of the Gal1P (**Fig. 6E**). In contrast, the representative complex extracted from local minimum of G1P reveals a differential induced fit conformation wherein there is change in hydrogen bond interactions involving the three hydroxyl groups. The 2OH group forms a hydrogen bond interaction with D93, 4OH forms a hydrogen bond interaction with D220, and the 6OH forms a hydrogen bond with H217. Consequently, there is a loss in the hydrogen bond interaction between 4OH and E213 (**Fig. 6F**). Collectively, these data further highlight that DdGal1Pase discriminates between substrates through an ensemble of subtle interactions, rather than a 1-2 key interaction.

**Fig. 6.**
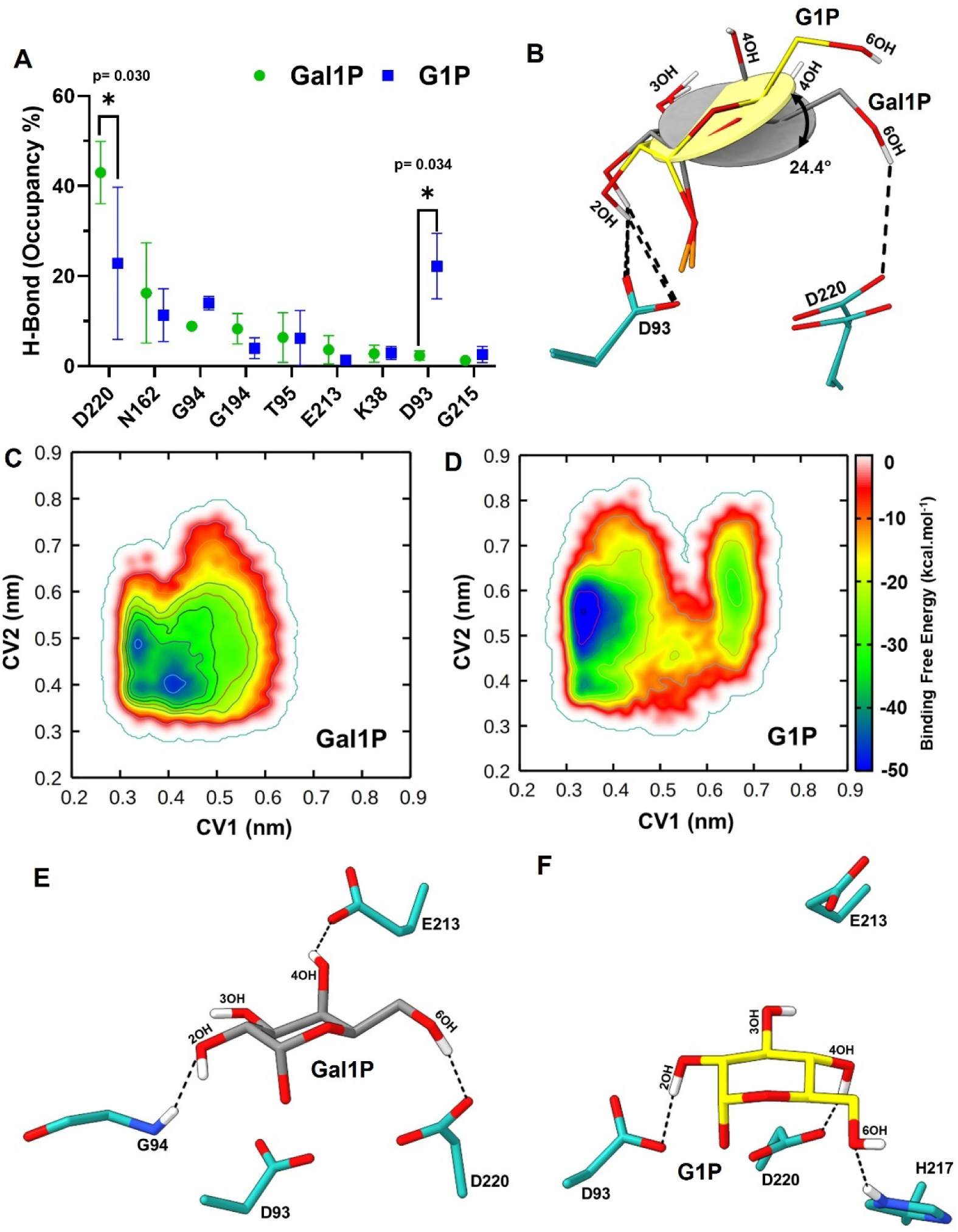
Simulations reveal selectivity of Ddgal1Pase towards Gal1P is governed by D93 and D220. (**A**) The average hydrogen bond occupancy between Gal1P or G1P and active site residues. Bars represent the mean ± SEM. p-values represent a two-way ANOVA. (**B**) The tilt angle of the pyranose ring of Gal1P (grey) and G1P (yellow) due to a change in occupancy of active site residues D93 and D220. The phosphate oxygens were hidden for clarity. Free energy landscape plots from well-tempered metadynamics for (**C**) Gal1P and (**D**) G1P. Representative structure depicting the local minima conformation of the (**E**) Gal1P (grey sticks) complex or (**F**) G1P (yellow sticks) complex. The phosphate group and the Mg^2+^ ions were hidden for clarity. The black dotted lines indicate the hydrogen bond interactions of the substrate with the labeled active site residues.

## DISCUSSION

In the past decade, research advances have enabled increasingly sophisticated methods for the synthesis of D-tagatose. While much progress has been made in improving the activity of isomerases and epimerases, these approaches often still suffer economic and efficiency challenges. Attempts to improve D-tagatose production schemes have generally focused on improving LAI activity, controlling conditions for more favorable equilibrium, or using alternative substrates (Baptista et al., 2021). Alternative approaches have made headway by exploring other routes through fructose epimerization, both in vitro (S. Liu et al., 2023) and in vivo (Dai et al., 2021), but need to overcome problems with substrate competition between fructose-6-phosphate and fructose 1,6-bisphosphate when implemented in whole cell contexts (Palur et al., 2025). Our study addresses these limitations by introducing a novel biosynthetic route that leverages whole-cell catalysis and a newly identified Gal1P-specific phosphatase to reverse the Leloir pathway. This strategy allows for direct conversion of glucose to galactose, providing a high-yielding precursor for tagatose production. Unlike prior strategies that rely on lactose-derived galactose and incur inherent 50 % substrate losses, our approach utilizes inexpensive glucose with a theoretical pathway yield of 94.9 %.

A key innovation of this work is the discovery and implementation of a highly selective Gal1P phosphatase from *Dictyostelium discoideum* (DdGal1Pase), which acts as the thermodynamic driving force to shift the equilibrium of the reversible Leloir pathway. Other inositol monophosphatases (IMPases) with substrate specificity to L-galactose-1-phosphate (Laing et al., 2004), or with promiscuous activity towards Gal1P (Yadav et al., 2020) have been described, but none that have shown the observed degree of specificity towards Gal1P demonstrated in this study. DdGal1Pase shares identity and a structurally conserved fold with many such characterized IMPases (**Fig.S23-S24)**, with the unique Gal1P specificity coming from nuanced active site residue variation. Consistent with previous reports, our hybrid QM/MM analysis supports the previously proposed two metal Mg² dependent mechanism, with the formation of a nucleophilic hydroxide from water to initiate catalysis (Bone et al., 1994; Ferruz et al., 2016; Wang and Hirao, 2013). As demonstrated through our modelling, DdGal1Pase’s active site is structurally optimized for Gal1P, enabling precise alignment of the sugar ring and phosphate group for catalysis. In contrast, the subtle stereochemical differences in G1P result in suboptimal steric and electrostatic complementarity within the active site, thereby reducing its catalytic efficiency. QM/MM reaction simulations show that Gal1P has a significantly lower activation free energy (TS1) than G1P (12.6 kcal·mol^−1^) for the phosphate hydrolysis reaction, implicating that Gal1P undergoes phosphate hydrolysis more readily than G1P, as in line with our experimental evidence. Furthermore, free-energy landscape analysis from well-tempered metadynamics reveals a pronounced minimum for the Gal1P-bound state, corresponding to a single, stable binding mode. By contrast, G1P produces a flatter free-energy landscape with multiple shallow minima, indicating it likely samples several weakly bound conformations.

We also demonstrated that the reverse Leloir pathway output is highly tunable and responsive to expression levels, especially of DdGal1Pase. This enzyme’s specificity for Gal1P over structurally similar phosphosugars is unprecedented and highlights a new axis for enzymatic control in sugar biosynthesis pathways. Without substantial optimization, the engineered *E. coli* strains produced up to 10.5 g/L galactose and > 1 g/L tagatose from 30 g/L glucose. These titers are promising for a first demonstration and suggest immediate industrial relevance with further strain and process improvements. Our analysis showed that tagatose yield was not limited by LAI activity under our conditions, but likely by the unfavorable equilibrium of the galactose-to-tagatose isomerization. We observed that increasing temperature after biomass accumulation improved this equilibrium, consistent with previous findings that LAI reactions at higher temperature favor tagatose formation at equilibrium (Bober and Nair, 2019). Given these results, it may be possible to improve tagatose yield further by increasing temperature with a thermostable LAI following the production of galactose, as well as by optimizing media conditions.

Additionally, engineering cellular export by deleting the sugar transporter gene *ydeA* improved intracellular tagatose levels, suggesting that transport bottlenecks can further be exploited to modulate pathway thermodynamics. This finding supports the notion that intracellular accumulation of galactose can shift the equilibrium toward tagatose, a concept that has previously been explored (Bober and Nair, 2019; Kim et al., 2008). Despite previous reports that MglB is the primary transporter responsible for import of galactose and export of tagatose (Kim et al., 2008), deletion or over-expression did not have an impact on tagatose production in our system. Further improvements in tagatose titers may be achieved by enhancing efflux capabilities. In *E. coli*, transport of sugars is regulated and is often limiting (Carreón-Rodríguez et al., 2023; Koita and Rao, 2012), suggesting that engineering relevant exporters could trap galactose and/or alleviate intracellular accumulation of tagatose. Furthermore, the yield of tagatose from glucose could be improved by reducing the amount of glucose fed to cells, likely due to a slower rate of galactose formation, and reduced loss of galactose to the extracellular medium. This could be exploited in fed-batch reactors by maintaining low glucose levels throughout the bioconversion.

Beyond its immediate application to tagatose, our pathway provides a generalizable framework for the biosynthesis of galactose-derived molecules directly from glucose. For example, redox-based tagatose pathways (Liu et al., 2019) or other galactose-dependent reactions could be paired with our system. The unique substrate specificity of DdGal1Pase could also provide a model for rational design of other phosphatases that could be used to generate free sugars like fructose for epimerization to tagatose (Shin et al., 2020), or in other rare sugar pathways such as allulose, which shares many of the same production challenges as tagatose (Guo et al., 2024). In summary, we demonstrate a platform to convert glucose to galactose and tagatose via Leloir pathway reversal. This was enabled by the discovery of a Gal1P-specific phosphatase and strategic pathway engineering. With further development, this system could support cost-effective, scalable production of tagatose and other galactose-derived products, reducing reliance on lactose and improving process sustainability.

## MATERIALS & METHODS

### Data Sources and Phylogenetic Analysis

Unless specifically noted, gene sequences were accessed from UniProt (Release 2025_03). Phylogenetic analysis was performed by performing multiple sequence alignment using MAFFT v7 with default parameters (Katoh and Standley, 2013). The resulting alignment was used to infer a maximum-likelihood phylogenetic tree using FastTree v2.1.11, employing the JTT+CAT model for amino acid substitution (Price et al., 2010). The resulting tree was visualized and annotated using FigTree v1.4.4 (http://tree.bio.ed.ac.uk/software/figtree/). Selected enzyme sequences were re-coded for *E. coli* expression using the codon optimization tool from Azenta Life Sciences (Boston, MA). See **Data S1** for sequences used in this study.

### In Vitro Enzyme Activity Assay

All in vitro enzyme expression was done using a pET-28a(+) vector. Sequences were ordered from Twist Biosciences (San Francisco, CA) and were designed to be cloned directly into a pET-28a(+) vector using an NcoI and HindIII double digest (New England Biolabs Ipswich, MA). Vectors were cloned using *E. coli* DH5α cells and sequence verified. Sequence verified pET-28a(+) vectors, as well as one additional control “blank” plasmid, which contained no cloned enzyme, were expressed in *E. coli* BL21(DE3) cells. Cells were grown in 50 mL tubes with 10 mL Lysogeny Broth (LB) medium + 50 µg/mL kanamycin at 37 °C for 16 h. The following day, cultures were diluted to OD = 0.05 in 50 mL tubes with 10mL LB + 50 µg/mL kanamycin and grown at 37 °C for 2 h. Cultures were then induced with 20 µL of 0.5 M IPTG (1 mM final concentration) and left to grow at 37 °C for an additional 4 h. to allow for protein expression. Subsequent steps were performed at 4 °C. Cells were chilled on ice for 20 min., and then centrifuged for 10 min. at 3200 × g. The supernatants were decanted, and cell pellets were resuspended in 5 mL of 0.25M HEPES, pH = 7. Samples were then sonicated with a Branson Sonifier 250 (Danbury, CT), with 5 s on 20 s off cycles for 24 cycles, using 40 % amplitude. Protein extracts were clarified by centrifugation at 20,000 × g. for 10 min. Aliquots were taken in microcentrifuge tubes and stored at –20 °C.

Cell-free extracts were evaluated for protein expression using a total protein Bradford assay (Pierce) and SDS-PAGE analysis. Cell-free extracts were screened with appropriate purchased pure substrates (> 95 %). Reactions were carried out in microcentrifuge tubes in 50 µL final volume using HEPES buffer (pH = 7). Generally, 30–60 µg/mL of total protein was loaded, and total protein concentration was normalized. Phosphatase reactions were supplemented with 1 mM MgSO_4_, while epimerase reactions did not contain any cofactors. All in vitro enzyme screens were controlled with a biological blank sample that consisted of cell-free extract from an empty pET28a(+) vector to control for background enzymatic activity. Reactions were initiated by substrate addition and allowed to incubate at 37 °C for 16 h. The reaction was stopped by the addition of 150 µL acetonitrile (75% final) and was analyzed using various LC methods.

### Plasmid and Strain Construction

Plasmid cloning was done by PCR amplification out of DNA templates followed by AarI mediated type IIs digestion and T4 ligase assembly. For all in vivo studies, an f1 origin bacterial artificial chromosome (BAC) was used. All cloned vectors were sequence verified by Sanger sequencing. Chromosomal integrations and chassis modifications were performed using the plasmid pKD46 encoding λ-Red recombinase machinery (Datsenko and Wanner, 2000). Edits were generally made using a positive selectable marker with a kanamycin resistance gene and the *sacB* gene encoding a levansucrase from *Bacillus subtilis*, which creates sucrose-dependent negative selection as previously described (Li et al., 2013). Deletions were made using the established method described by Datsenko and Wanner (Datsenko and Wanner, 2000). All chromosomal edits were verified by PCR of the modified locus and sanger sequencing. Genetic deletions indicated with “Δ” were designed to remove the entire coding sequence of the indicated gene. Chromosomal integrations replaced the native gene coding sequence with the recombinant expression cassette. Integrations into intergenic sites denoted by “int(gene1-gene2)” splice in the DNA expression cassette between the two indicated genes in the *E. coli* chromosome.

### Tagatose Production Assays

Strains of *E. coli* were transformed with BAC plasmids using a BioRad Genepulser Xcell electroporator (Hercules, CA) with 2.5 kV and then recovered in SOC media for 1 h. The SOC recoveries were then plated onto LB agar plates containing chloramphenicol as a selection antibiotic. For chromosomal variants, strains were streaked out directly from glycerol stocks onto LB agar. The plates were grown overnight in a stationary incubator at 37 °C. The following day, colonies from the transformations were seeded into 96 deep well plates (DWPs) containing 400 µL of rich fermentation growth media containing either galactose or glucose. Briefly, the phosphate-buffered nitrogen-limited media contained 5 g/L yeast extract (Nucel), ammonium chloride, ammonium sulfate, magnesium sulfate, citric acid, ferrous sulfate, thiamine HCl, and standard trace metals. The wells were seeded in triplicate, with a single colony used to inoculate each well. Seed cultures were incubated at 37 °C in a 1-in orbital shaking incubator at 250 rpm for 24 h. Production cultures were inoculated from the 24 h seed cultures by adding 40 µL of the seed culture to 360 µL of fresh fermentation growth media in a 96DWP. Production cultures were incubated at 37 °C in a shaking incubator at 250 RPM for 48 h before analysis by LC.

### Analytical Methods

For in vivo assays, liquid-liquid extractions were done for the purpose of extracting reaction products and intermediates from the cell cultures into a matrix that could be analyzed by the applicable analytical instrumentation. The extraction was performed in situ by adding 400 µL (equal volume) of 100 % acetonitrile, pulse vortexing for 10 min, and centrifuging at 3200 × g for 10 min. Supernatants were diluted an additional 20–50× depending on how much glucose or galactose was fed. 75 % acetonitrile was used for LC/MS dilutions, and pure water was used for UPLC-RID dilutions. Diluted samples were run by LC/MS for nucleotide sugars, sugar phosphates, and tagatose quantification, and on the UPLC-RID for glucose and galactose quantification.

Quantitative analysis of free glucose and galactose by UPLC-RID was performed on an Agilent 1260 Infinity II UPLC with a Refractive Index Detector (RID). The column used was a Rezex RCM-Monosaccharide Ca^2+^, 100 × 7.8 mm controlled at 80 °C. The method was isocratic with 100% water with a 15-min run time. Tagatose was measured separately using a Milipore SeQuant ZIC-pHILIC 150 × 4.6 mm column and an 18-min gradient between solutions of aqueous 100 mM ammonium bicarbonate and 100 % acetonitrile. Detection was done using electrospray ionization with an Agilent 6460 triple quadrupole mass spectrometer and monitoring a selected ion for tagatose.

For in vitro epimerase enzyme discovery workflows, analysis of UDP-galactose, UDP-glucose, galactose-1-phosphate, and glucose-1-phosphate was performed using a ThermoFisher Scientific Accucore-150mm Amide HILIC column (16726-152130). This method used an 81-min gradient between mixtures of aqueous 100 mM ammonium acetate pH 9.5 and 100 % acetonitrile. For in vitro phosphatase analysis, galactose-1-phosphate, glucose-6-phosphate, glucose-1-phosphate, galactose, and glucose were detected using a Milipore SeQuant ZIC-pHILIC 150 × 4.6 mm column with a 31-min gradient between solutions of aqueous 100 mM ammonium bicarbonate and 100 % acetonitrile. Detection for both methods was done using electrospray ionization with an Agilent 6460 triple quadrupole mass spectrometer and monitoring selected ions from each species.

### Data Analysis

All data were collected from distinct samples. Mean, standard deviation, and general statistics were calculated using built-in functions in Matlab® or Microsoft Excel. P values < 0.05 are generally considered statistically significant. Unless otherwise stated, error bars in all plots represent standard deviation.

### Structural modelling of DdGal1Pase and deriving the plausible enzyme substrate complexes for Gal1P and G1P

The DdGal1pase structure (UNIPROT ID: Q54U72) from *Dictyostelium discoideum* was modelled using AlphaFold3 (Abramson et al., 2024). The stereochemical quality of the modelled structure was assessed using the Structure Assessment Webserver (Waterhouse et al., 2024) (https://swissmodel.expasy.org/assess). Placement of Mg^2+^ ions was based on the position of Mn^2+^ ions in the human inositol monophosphatase structure (PDB ID: 1IMC) (Bone et al., 1994). As DdGal1Pase was identified as an Mg^2+^-dependent enzyme, three Mg ions were modelled in the *apo* form of DdGal1Pase. For deriving the DdGal1Pase substrate complex with Gal1P and GlP, _3_Mg^2+^ was removed and only the catalytically essential _2_Mg^2+^, and _1_Mg^2+^ were retained (Atack et al., 1995; Bone et al., 1994; Wang and Hirao, 2013). Molecular docking of Gal1P and G1P were performed using Autodock4.2 (Morris et al., 2009) to derive the plausible enzyme substrate complexes. The Lamarckian genetic algorithm (LGA) was applied to determine the most plausible enzyme substrate complexes based on the previous reported interactions of inositol (Atack et al., 1995) and based on the catalytic mechanism of *myo*-inositol monophosphatase (Wang and Hirao, 2013). A grid of 60, 60, and 60 points in the *x, y*, and *z* directions, respectively, with a grid spacing of 0.375 Å was used around the active site pocket. Grids were prepared to reflect the areas of the catalytic residue (E71) and active site residues (D90, I92, D93, G94, T95, A196, E213 and D220). The number of poses per substrate that passed the grid refinement calculation was set to 500. All substrates were drawn using MarvinSketch, and structures were minimized using the Universal force field (UFF) (Rappe et al., 1992) in Avogadro (Hanwell et al., 2012). University of California, San Francisco (UCSF) ChimeraX was used for visualization and structural figures were also generated using the same tool (Goddard et al., 2018; Meng et al., 2023; Pettersen et al., 2021).

### Molecular dynamics and metadynamics simulation parameters for DdGal1Pase-substrate complexes

The molecular dynamics (MD) simulations of DdGal1Pase-Gal1P, DdGal1Pase-G1P complex, and DdGal1Pase (*apo*) were simulated using Amber99sb force field (Lindorff Larsen et al., 2010) as implemented in GROMACS 2023 (Abraham et al., 2015). The DdGal1Pase built complex systems were solvated using the TIP3P water model (Jorgensen et al., 1983) and neutralized with salts ([NaCl] = 0.15 M). The Particle Mesh Ewald (PME) technique (Darden et al., 1993) was utilized to compute electrostatics using a real-space cut-off of 10 Å. Lennard-Jones 6-12 potentials with a 14 Å cut-off were used to represent Van der Waals interactions. Linear Constraint Solver (LINCS) (Hess et al., 1997) was used to limit hydrogen bonding, while V-rescale was used to maintain the temperature at 300 K. Using the steepest descent approach, energy minimization was performed to obtain a maximum force of 10 kJ/mol. For all molecular dynamic simulations, the time step was set to 2 fs. The minimized systems were subjected to NVT (canonical ensemble) and NPT (isothermal-isobaric ensemble) equilibration for 1 ns each at 300 K prior to the production run. The partial charges for the substrates and products were derived using density functional theory at the B3LYP/6-311++ G (2d, p) level in the water system and implemented using GAMESS version 30 (Barca et al., 2020). The substrate and product force-field parameters were estimated using ACPYPE (Sousa da Silva and Vranken, 2012) and the ANTECHAMBER module of AMBER11 (Case et al., 2023). The Gal1P and G1P substrates were simulated in ionized forms with -2 charges. The entire process of simulating the substrates and *apo* was carried out three times with 200 ns each complex, totaling 600 ns of simulations. All six systems achieved convergence within 200 ns of the simulation, totaling 1.8 µs of simulation time with an average number of atoms per system of ∼23,552 atoms across various simulations. A system with varying compositions of atoms and their specifics is presented in **Table S3**. The hydrogen bond occupancy was analyzed using HBonds plugin, v1.2 from VMD (Humphrey et al., 1996). Furthermore, a two-way ANOVA statistical analysis was conducted, followed by a Sidak’s correction to control type I statistical errors arising after the ANOVA analysis. The significance was kept at 95%.

Well-tempered meta dynamics (MTD) simulations (Laio and Parrinello, 2002) were conducted utilizing the PLUMED library (Bonomi et al., 2009), employing structures obtained after 1 ns of NPT simulation. Metadynamics is an enhanced sampling method that calculates the free energy surface along predefined collective variables (CVs) by applying a Gaussian bias potential along the CVs as they are sampled during the simulation. Well-tempered MTD also employs a factor to decrease the magnitude of the bias potential at CV values that are heavily sampled, thereby ensuring rapid convergence. In this study, two distance-based collective variables (CVs) were selected based on hydrogen occupancy analysis from conventional MD simulations. The CVs were selected where in CV1 is the distance between the center-of-mass (COM) of D93 (CB, CG, OD1, and OD2 atoms) and the C2OH group of Gal1P/G1P, while the second collective variable (CV2) represents the distance between the center-of-mass (COM) of D220 (CB, CG, OD1, and OD2 atoms) and the C6OH group of Gal1P/G1P. The DdGal1Pase-Gal1P and DdGal1Pase-G1P complexes were simulated for 2 × 20 ns for each complex, utilizing different seeds for initial velocity generation, with PACE = 500, HEIGHT = 1.0, SIGMA = 0.01, bias factor of 2, and temperature of 310 K.

### QM/MM Simulations Parameters of DdGal1Pase-substrate Complexes

The DdGal1Pase-substrate complexes (Gal1P and G1P) were initially prepared by embedding them in a cubic box with a 12 Å solute-solvent separation margin in all dimensions, using the QwikMD (Barreto Gomes et al., 2022) program integrated into VMD (Humphrey et al., 1996). To maintain electroneutrality, Na^+^ & Cl ions were added to achieve a salt concentration of 0.15 M. The CHARMM36 (Huang and MacKerell, 2013) force field was employed to parameterize the protein topology using the psfgen and autopsf modules. The Gal1P and G1P were parameterized using CGenFF web server (Vanommeslaeghe et al., 2010) (https://cgenff.com/).

The system solvation was handled using the TIP3P (Mark and Nilsson, 2001) water model. Short-range non-bonded interactions were treated with a 12.0 Å cutoff, while long-range electrostatics were computed using the Particle-Mesh Ewald (PME) method. The r-RESPA multiple time step scheme was applied, with short-range interactions updated every step and long-range interactions every two steps, using a 2-fs integration time step. System equilibration began with a 1000-step energy minimization using the conjugate gradients method, followed by heating to 300 K via the Langevin thermostat (collision coefficient of 1 ps ¹). The pressure was maintained at 1 atm using a barostat. For QM/MM simulations, the QM region was defined to include E71, E72, D48, D90, I92, D93, D220, Gal1P/G1P, Mg ions and modelled three water molecules with the total number of atoms 107 in the QM region. The charge of each QM region was kept between +1 and −1 to ensure effective semi-empirical QM calculations. Subsequent steps included 1000 steps of minimization and 10,000 steps of simulated annealing for equilibration. The equilibrated QM/MM structure then served as the initial input for QM/MM simulations. The simulations were performed at 300 K and 1 bar under periodic boundary conditions, with a 0.5-fs time step ensuring stability over 1000 ps of simulation time. The production phase of the hybrid QM/MM simulations was conducted using the MOPAC (Stewart Computational Chemistry - MOPAC) PM7 semi-empirical QM method in conjunction with the CHARMM36 force field (Huang and MacKerell, 2013), ensuring that the total charge remained between +1 and -1. The PM7 semi-empirical method was employed for geometry optimization and electronic structure calculations, with a cutoff of 9.0 Å for nonbonded interactions with incorporating single-point SCF calculations and gradient evaluations for system relaxation. All the QM/MM simulations were performed using NAMD (Phillips et al., 2020).

## AUTHOR CONTRIBUTIONS

A.M.L. and C.G.T. conceptualized the study, performed the experimental work, acquired and curated the data, carried out formal analysis and data visualization, and wrote/edited the manuscript. H.L.L. performed the transporter study. P.K.A. edited the manuscript. N.U.N. helped conceptualize the study, plan the experimental methodology, and edit the manuscript. C.N.S. helped edit the manuscript and provided project administration and supervision. P.K, A.K, S.N.K, and D.K.S were involved in technical discussions and in designing all the in silico experiments. A.K, S.N.K, DK and L.M performed all the in silico experiments. A.K, S.N.K, D.K and P.K contributed to writing the in silico section, reviewing the manuscript, and making revisions.

## DATA AND MATERIALS AVAILABILITY

All data needed to evaluate the conclusions in the paper are present in the paper and/or the Supplementary Materials.

