## Supplemental Information for "A Novel Phosphatase Reverses the Leloir Pathway to Promote Tagatose Synthesis from Glucose"

**GLOSSARY**

**Glucose:** D-glucose

**Galactose:** D-galactose

**Tagatose:** D-tagatose

**UDP:** Uridine diphosphate

**G6P:** D-glucose-6-phosphate

**G1P:** D-glucose-1-phosphate

**Gal6P:** D-galactose-6-phosphate

**Gal1P:** D-galactose-1-phosphate

**Tag6P:** D-tagatose-6-phospahte

**Gal1Pase:** D-galactose-1-phosphatase

**DdGal1Pase:** D-galactose-1-phosphatase from *Dictyostelium discoideum*

**LAI:** L-arabinose isomerase

**BcLAI:** L-arabinose isomerase from *Bacillus coagulans*

**pTM:** Predicted template modelling score

**pLDDT:** Predicted local distance difference test

**MD:** Molecular dynamics

**RMSD:** Root mean square deviation

**TS1:** Transition state 1

**INT1:** Intermediate state 1

**CV:** Collective Variable

**COM:** Center of mass

**IMPase:** Inositol monophosphatase

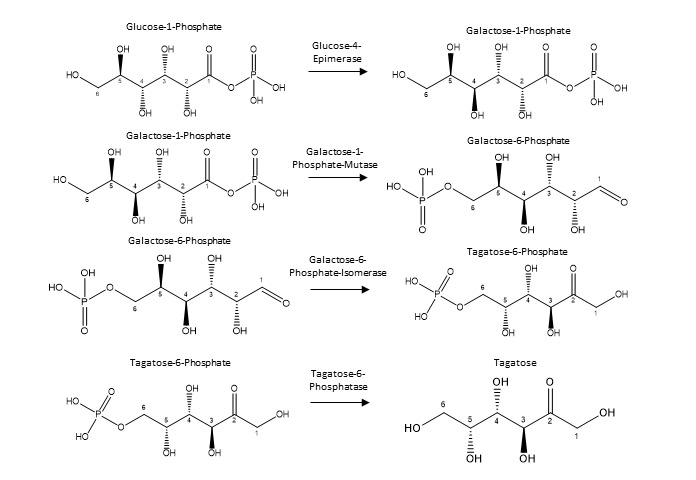

**Fig. S1. Production of tagatose from G1P using novel proposed epimerase and mutase reactions.** G1P is naturally produced in organisms like *E. coli* by phosphoglucomutase (*pgm*) from G6P. A novel glucose-4-epimerase and galactose-1-phophate-mutase could yield Gal6P from glucose, which could then be converted to Tag6P via the naturally occurring Gal6P isomerase. A terminal phosphatase which has also been previously demonstrated could then convert Tag6P to tagatose.

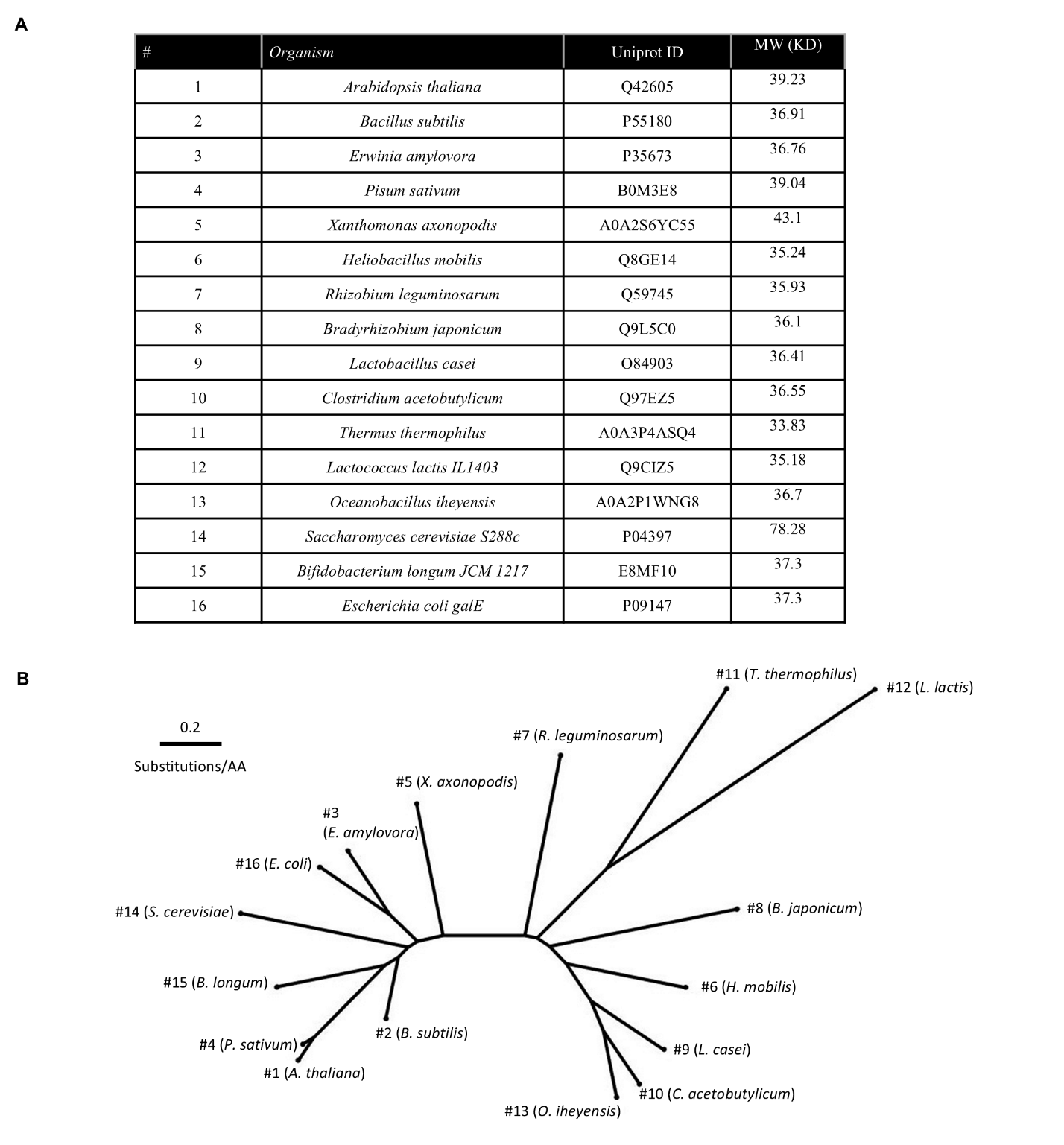

**Fig. S2. Epimerases tested for glucose-1-phosphate to galactose-1-phosphate activity.** (**A**) Table of 16 epimerases tested from a diverse set of organisms. (**B**) Phylogenetic tree representing 16 tested epimerases

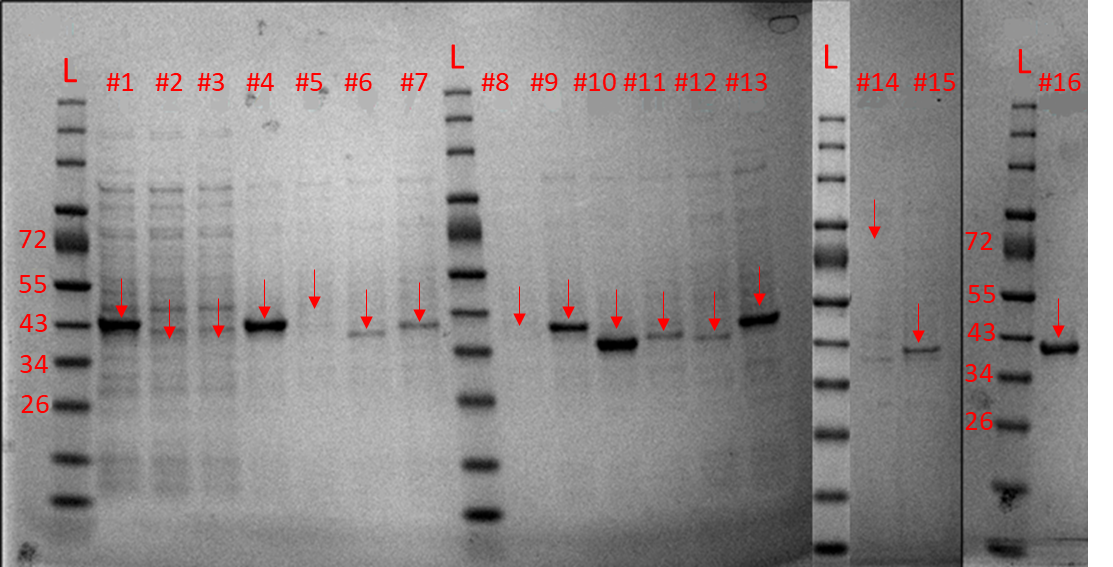

**Fig. S3. SDS PAGE of cell free extracts used in epimerase assays.** Ladder (L) represents size in kDa. Arrows represent the expected region where the expressed protein should appear. The figure has multiple gels combined together, the closest ladder to the left should be used to approximate protein size. Epimerases #2, #3, #5, #8, and #14 were removed from further analysis due to their lack of expression. Refer to Fig. S2 for enzyme numbering.

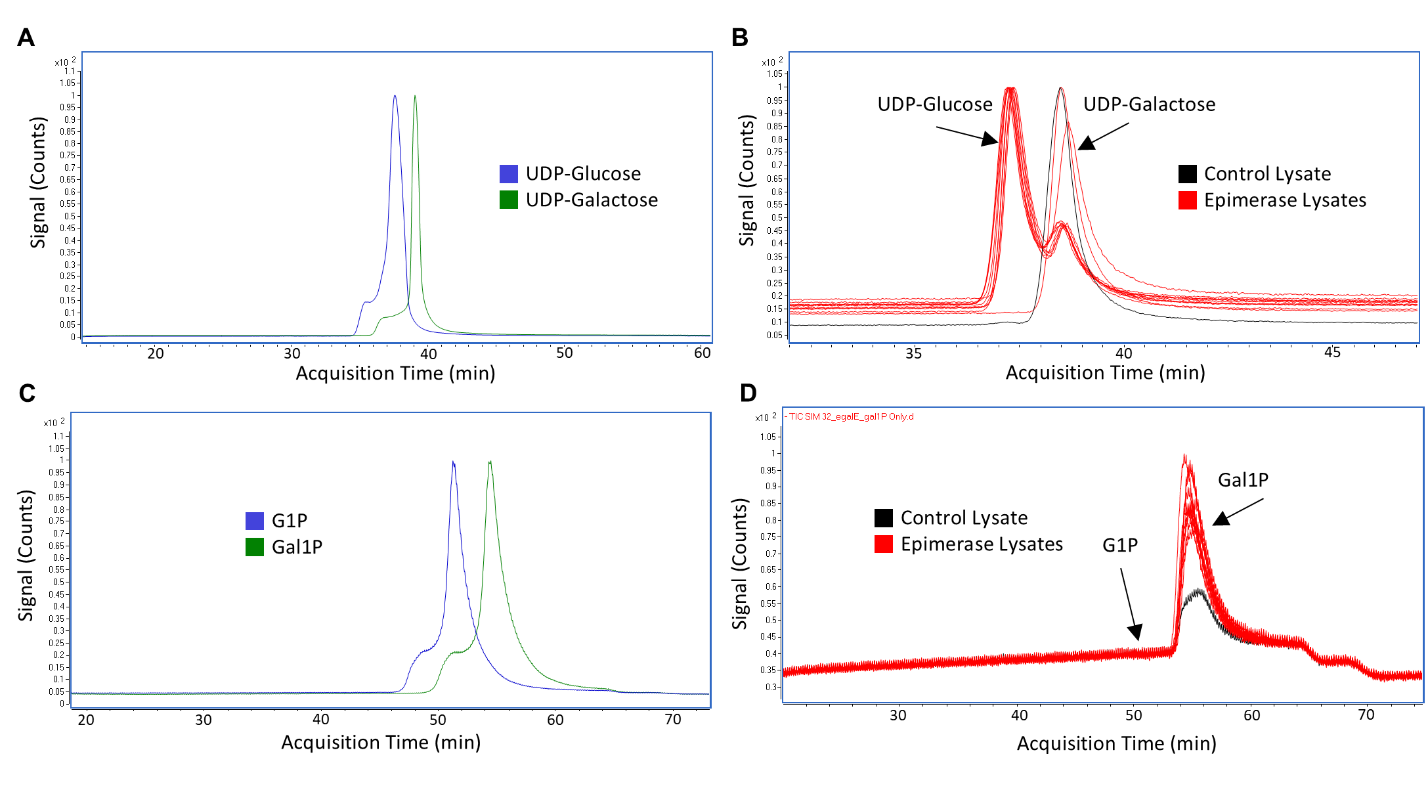

**Fig. S4. In vitro epimerase assay data.** The epimerases expressed in cell free extracts were tested with fed (**A and B**) UDP-galactose or (**C and D**) Gal1P and analyzed by LC/MS. Epimerases #1, #4, #6, #7, #9, #10, #11, #12, #13, #15, and #16 were screened for activity. Cell free extract without any over-expressed enzyme was sued as control. Refer to Fig. S2 for enzyme numbering. (**A**) UDP-galactose and UDP-glucose standards, as well as (**C**) G1P and Gal1P standards could be separated using an Amide HILIC column. Cell free extracts were tested for native substrate activity by feeding UDP-galactose and monitoring formation of UDP-glucose. All epimerases converted the majority of UDP-galactose to UDP-glucose except for #1 (lower activity), and #12 (no activity). The *E. coli* background control showed negligible activity. (**D**) Despite showing high activity on fed UDP-galactose, we did not observe any detectable activity for the interconversion between Gal1P and G1P.

**Glycolysis:**

$$Glucose+{2 NAD}^{+}+2 ADP+ {2 P}_{i}$$

$$\boldsymbol{\to}2 Pyruvate+2 NADH+{2 H}^{+}+2 ATP+{2H}_{2}O$$

**Pyruvate Dehydrogenase:**

$$2 Pyruvate+{2 NAD}^{+}+2 CoASH$$

$$\to2 \text{acetyl-CoA}+2 NADH+{2 CO}_{2}$$

**TCA Cycle:**

$$2 \text{acetyl-CoA}+{4 NAD}^{+}+{2 NADP}^{+}+ 2Q8+2 ADP+ {2 P}_{i}$$

$$\to2 CoASH+4 NADH+2 NADPH+2Q8H_{2}+2 ATP+{4 CO}_{2}$$

**Respiration:**

$$10 NADH+{10 H}^{+}+{5 O}_{2}\to{10 H}_{2}O+{10 NAD}^{+}+13.33 ATP$$

$$2Q8H_{2}+O_{2}\to{2 H}_{2}O+1.33 ATP+2 Q8$$

Net ATP:

| Glycolysis | 2 |
| --- | --- |
| TCA Cycle | 2 |
| Respiration | 14.67 |

**Net equation for respiration:**

$$Glucose+ {6 O}_{2}+18.67 ADP\to{6 CO}_{2}+{6H}_{2}O+18.67 ATP$$

**Net equation for tagatose pathway:**

$$18.67 Glucose+18.67 ATP\to18.67 Tagatose+18.67 ADP$$

**Combined equation for tagatose production:**

$$19.67 Glucose+ {6 O}_{2}\to{6 CO}_{2}+{6H}_{2}O+18.67 Tagatose$$

**Theoretical pathway yield:**

$$19.67 Glucose \times180.16\frac{g}{\mathrm{mol}}\to18.67 Tagatose\times180.16\frac{g}{\mathrm{mol}}$$

$$\boldsymbol{18.67\div19.67=94.9\%}$$

**Fig. S5. Theoretical yield for production of D-tagatose** **using the reverse Leloir pathway.** Basic equations are written for net cofactor production from glycolysis, pyruvate dehydrogenase, the TCA cycle, and cellular respiration. In this case, the only theoretical pathway yield loss comes from glucose used to make ATP for transport and phosphorylation of glucose to G6P. Calculations assume a transhydrogenase converts excess NADPH to NADH, a P/O ratio of 0.67 for ubiquinol-8 (Q8H_2_), and a P/O ration of 1.33 for NADH based on published values (Varma and Palsson, 1993). Complete oxidation of a single glucose is conservatively estimated to yield 18.67 ATP.

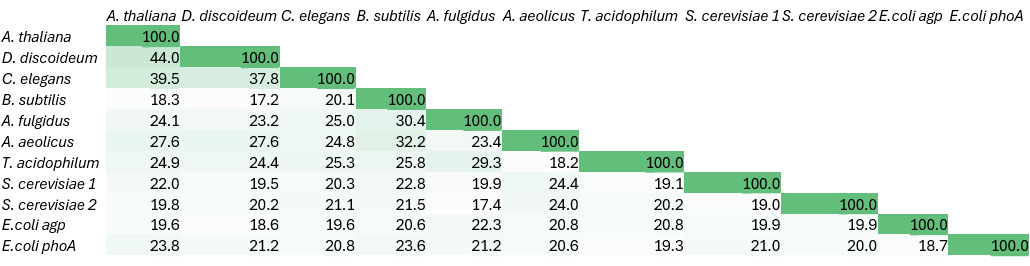

**Fig. S6. Pairwise sequence ID between phosphatases screened.** Amino acid sequences (**Data S1**) for 11 phosphatases screened for activity were compared and demonstrate low pairwise sequence identity. Global sequence alignments were performed using a Needleman-Wunsch algorithm.

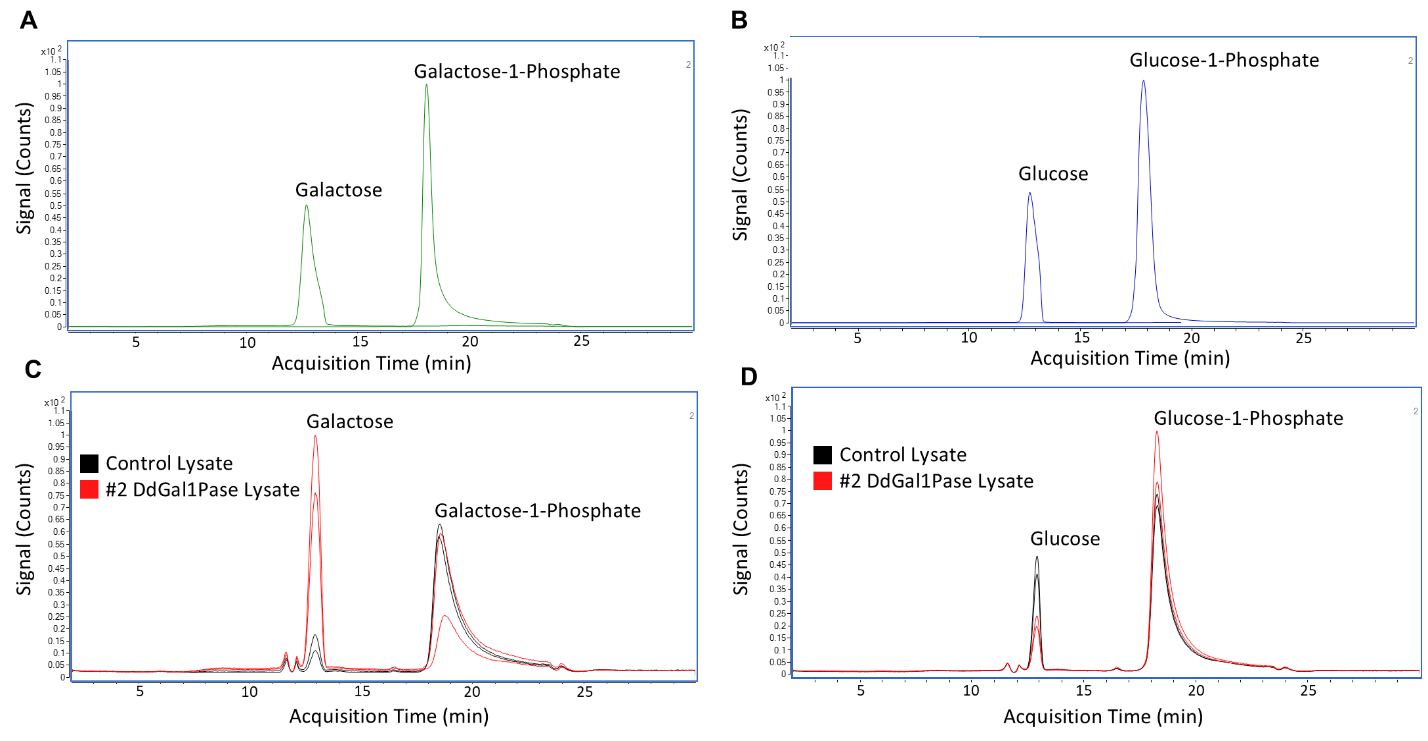
**Fig. S7. DdGal1Pase exhibits substrate preference for galactose-1-phosphate over glucose-1-phosphate in vitro.** Standards for (**A**) galactose and galactose-1-phosphate, or (**B**) glucose and glucose-1-phosphate, were separated and detected using LC/MS with a HILIC column. DdGal1Pase shown in **Figure 2** expressed in an *E. coli* cell free extract was tested with fed galactose-1-phosphate and glucose-1-phosphate to analyze enzymatic specificity. The cell free extract with DdGal1Pase over-expression demonstrates substantial activity on (**C**) galactose-1-phosphate over the *E. coli* background lysate control but (**D**) does not show activity over background when fed the C4 epimer glucose-1-phosphate.

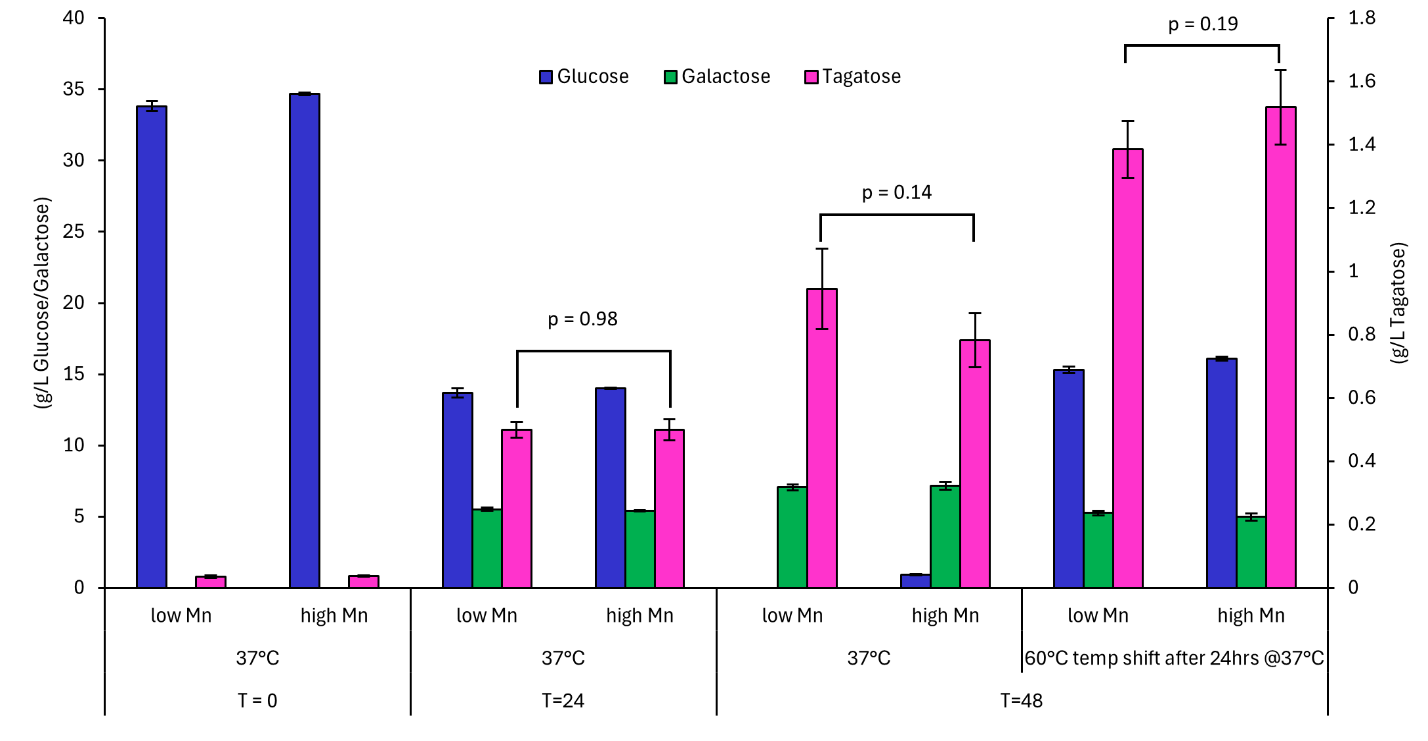

**Fig. S8. Manganese supplementation does not improve tagatose formation**. We evaluated performance from our fully integrated tagatose pathway strain (Δ*galKM*, Δ*pgi*::*pgm*, Δint(*rsmG*-*atpI*)::*BcLAI*, Δint(*lafU*-*dinB*)::*DdGal1Pase*) with or without a temperature shift to 60 °C starting at 24 h, and with or without Mn^2+^ supplementation to promote LAI activity. While shifting to a higher temperature promotes conversion from galactose to tagatose, increasing manganese from 0.05 mM to 0.45 mM did not facilitate significant improvement in tagatose formation. Bars represent means ± SD, n = 3.

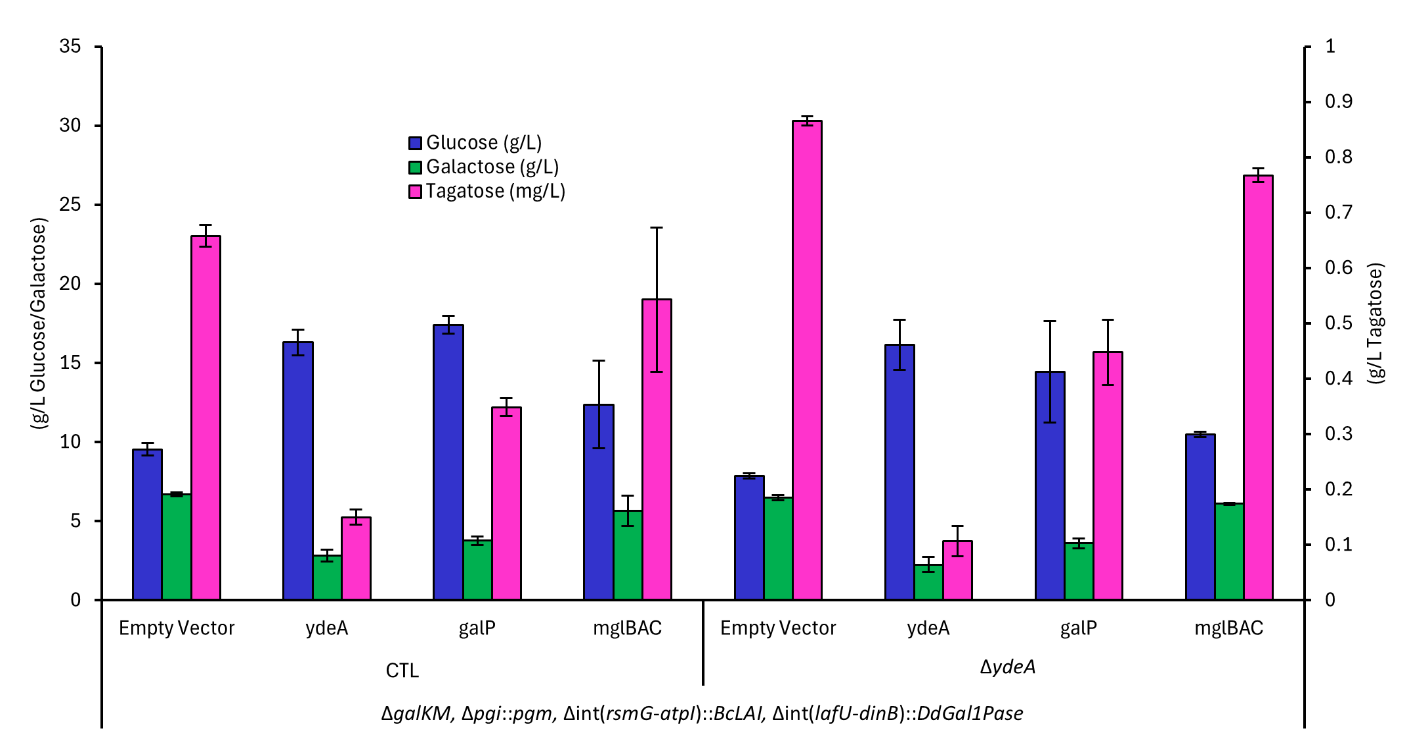

**Fig. S9. Transporter over-expression in tagatose production strain.** Select transporters were overexpressed using a weak constitutive promoter in the fully integrated tagatose pathway strain (Δ*galKM*, Δ*pgi*::*pgm*, Δint(*rsmG*-*atpI*)::*BcLAI*, Δint(*lafU*-*dinB*)::*DdGal1Pase*). Overexpression of select transporters generally reduces tagatose formation but also appears to inhibit glucose consumption. Bars represent means ± SD, n = 3.

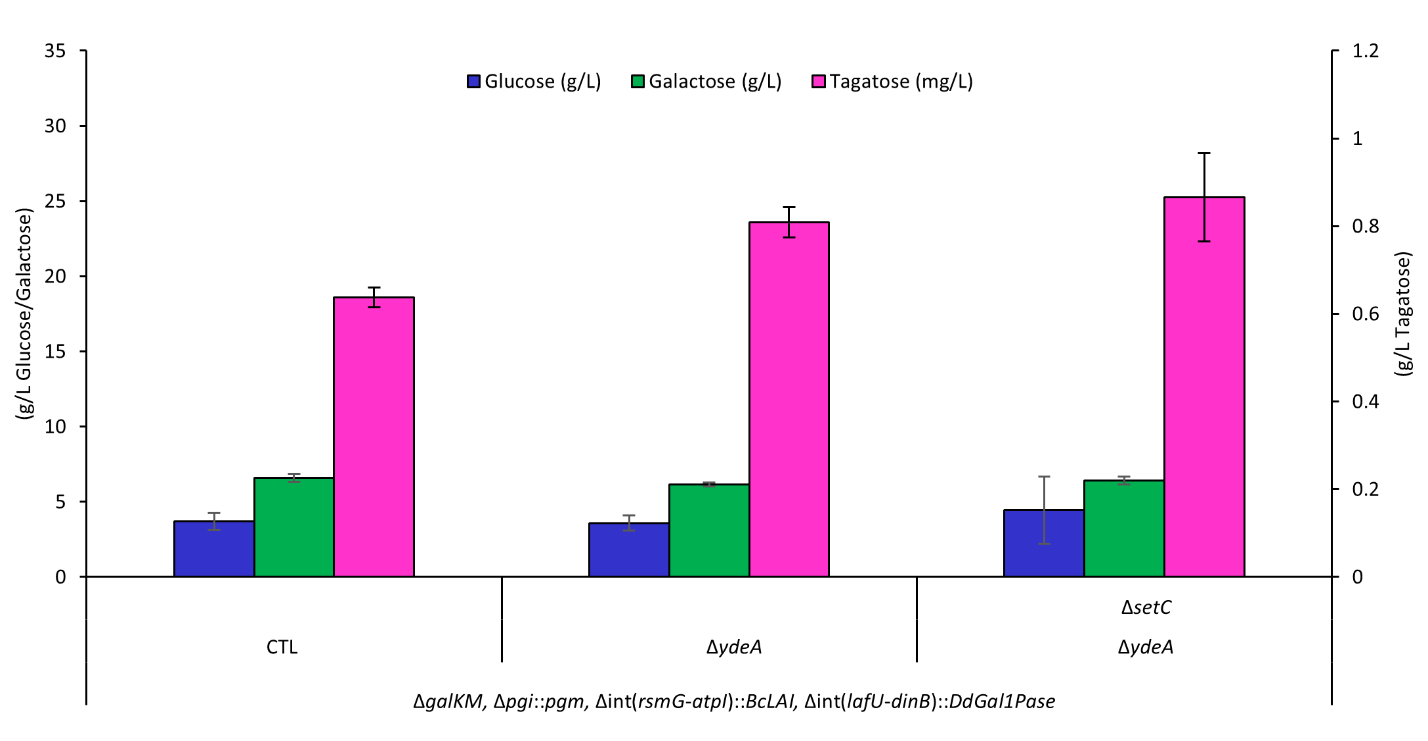

**Fig. S10. Transporter knockout combinations.** The fully integrated tagatose pathway strain in the fully integrated tagatose pathway strain (Δ*galKM*, Δ*pgi*::*pgm*, Δint(*rsmG*-*atpI*)::*BcLAI*, Δint(*lafU*-*dinB*)::*DdGal1Pase*) was further modified with multiple transporter deletions. While *ΔydeA* is beneficial, combining *ΔydeA* with Δ*setC* does not further improve tagatose formation. Bars represent means ± SD, n = 3.

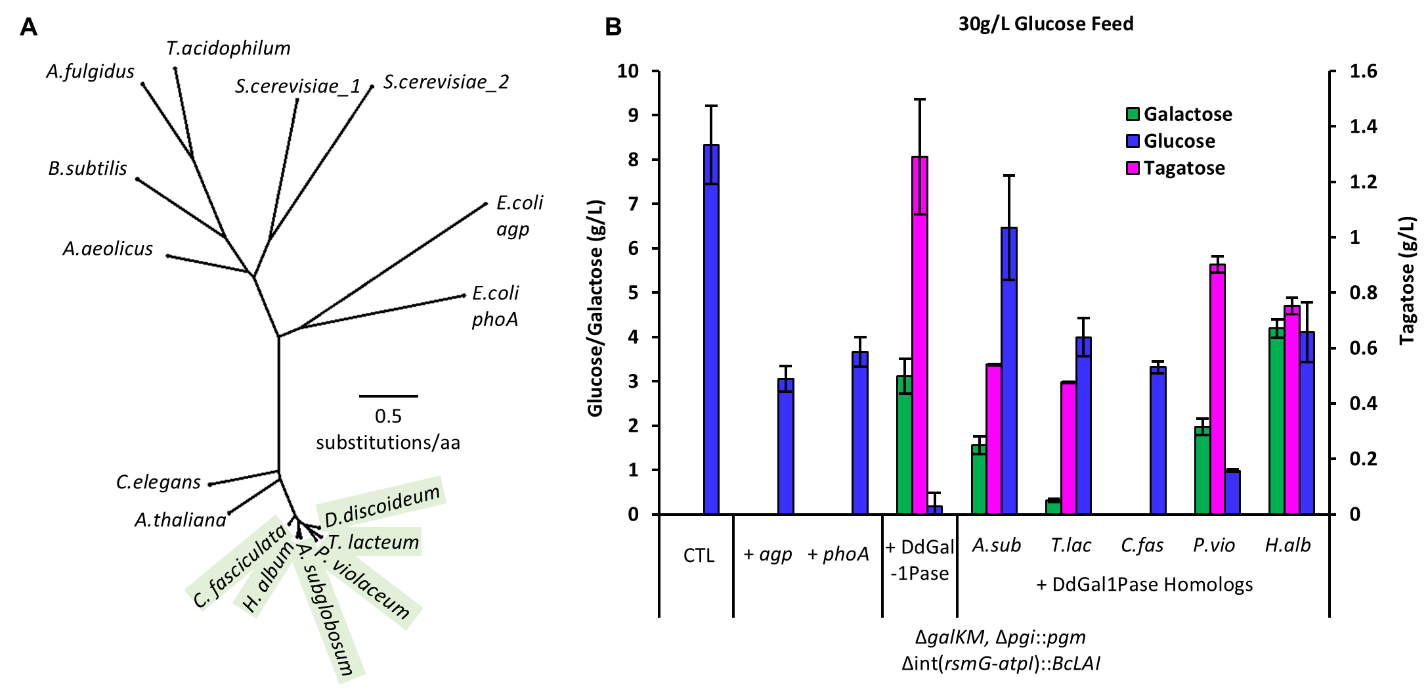

**Fig. S11.** Galactose-1-Phosphatases are phylogenetically and structurally specific. (**A**) Phylogenetic tree representing 5 additional sugar-phosphatase homologs (highlighted) tested with sequence similarity to DdGal1Pase in context with other phosphatases previously tested. (**B**) Galactose, tagatose, and glucose titer results from complementing DdGal1Pase, 5 homologs, or two native *E. coli* phosphatases into the *ΔgalKM Δpgi::pgm BcLAI* strain. 4/5 homologs enable Leloir pathway reversal, while native *E. coli* (nonspecific) phosphatases do not. Bars represent means ± SD, n = 3.

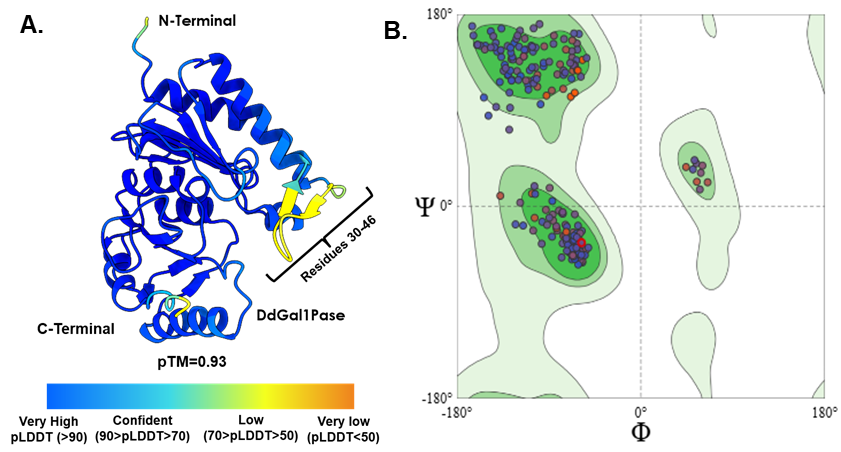

**Fig. S12. Validating structural model for DdGal1Pase. (A**) The AlphaFold modelled structure of DdGal1Pase with pLDDT confidence score. (**B**) Ramachandran plot for the modeled structure, where 97.41 % of the residues were in favorable regions.

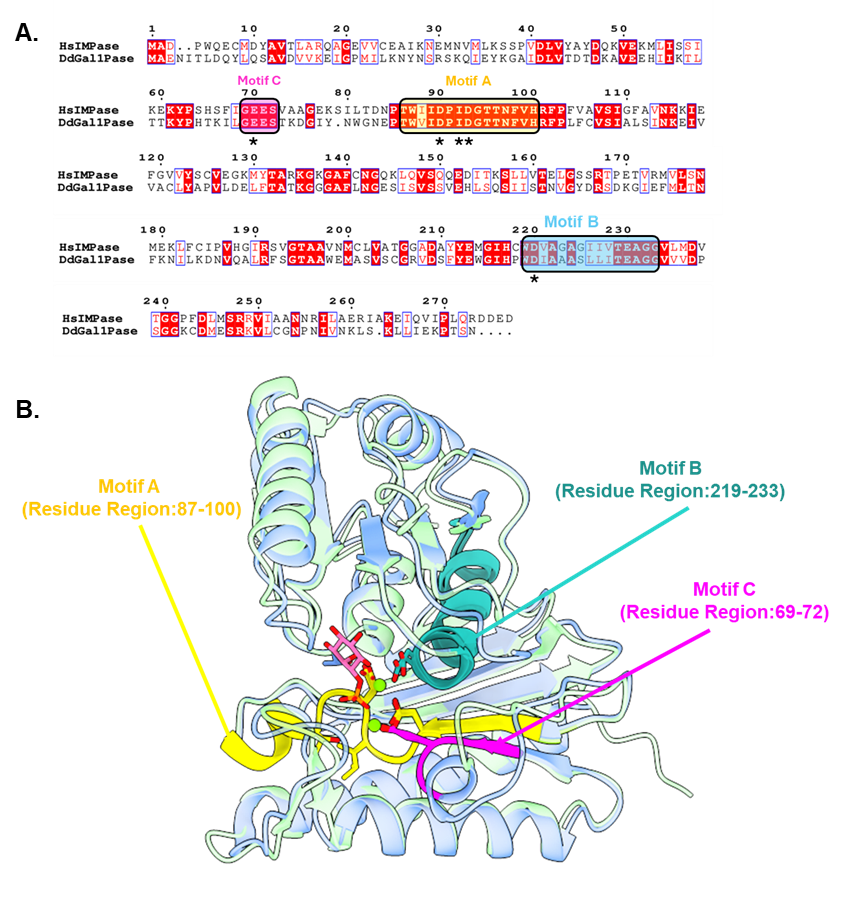

**Fig. S13.** (**A**) Multiple sequence alignment of human IMPase and DdGal1Pase, with identified structural motifs highlighted in boxes. The asterisks indicate ion-binding residues. (**B**) Structural superimposition of the crystal structure for human IMPase (PDB ID: 1IMA) (blue) and AlphaFold modeled DdGal1Pase (green) with motifs A, B, and C shown in yellow, teal, and magenta, respectively. The substrate inositol-1-phosphate is shown as pink sticks, and the two catalytic Mg^2+^ ions as green spheres.

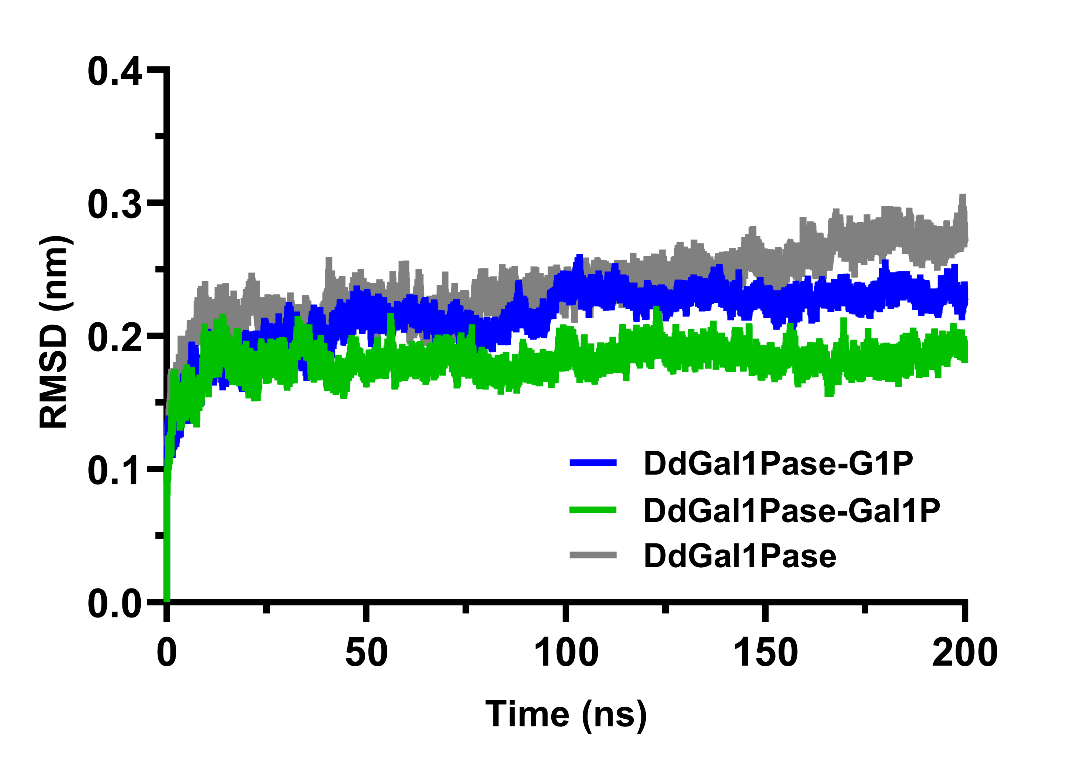

**Fig. S14.** Root mean square deviation (RMSD) of the DdGal1Pase backbone without and with substrate (G1P or Gal1P). Gal1P improves enzyme stability and promotes a more catalytically competent state. The plotted RMSD is the mean of 3×200 ns runs. G1P = glucose-1-phosphate, Gal1P = galactose-1-phosphate.

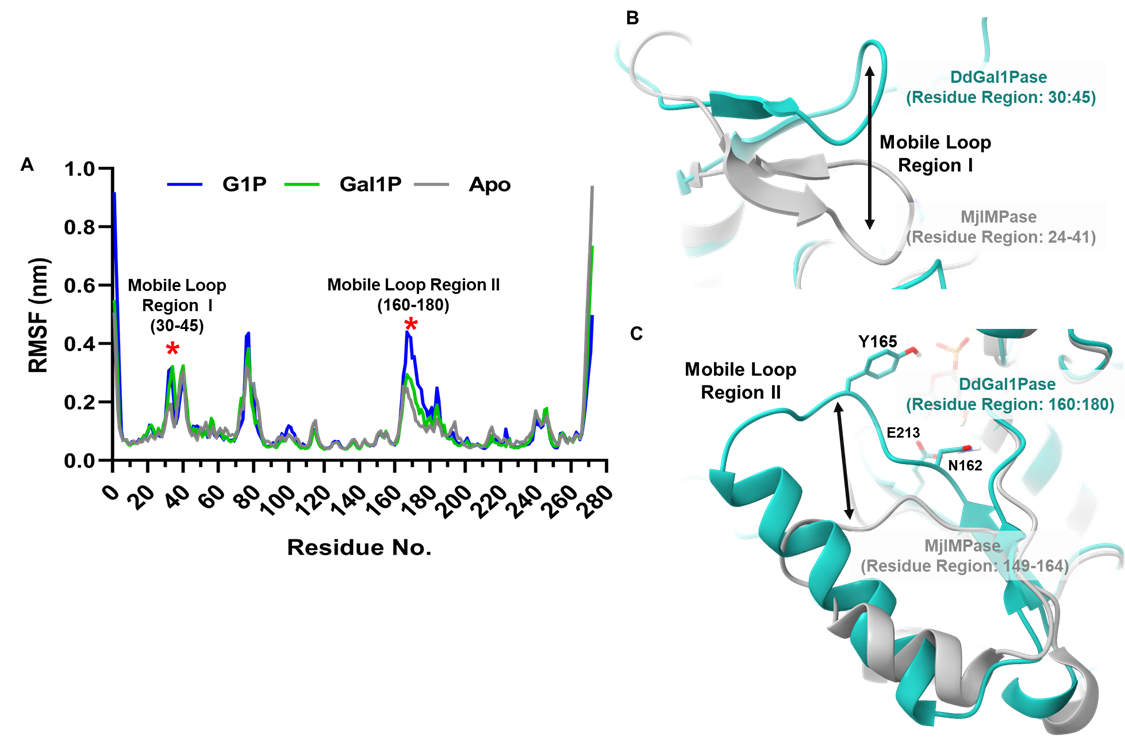

**Fig. S15.** (**A**) The root mean square fluctuation (RMSF) of DdGal1Pase and it’s substrate complexes, along with the identified mobile loops. (**B**) The mobile loop region I (MLR I) of DdGal1Pase and MjIMPase. (**C**) The mobile loop region 2 (MLR II) of DdGal1Pase and MjIMPase. MjIMPase is an inositol monophosphatase from *Methanocaldococcus jannaschii* (PDB ID: 1DK4).

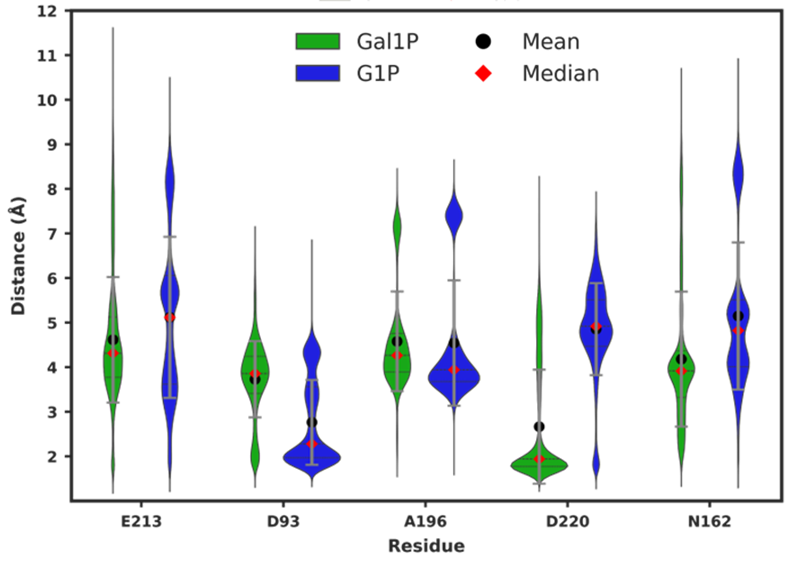

**Fig. S16.** Violin plot of the hydrogen bond distances of substrate pyranose ring hydroxyl groups with the active site residues; The vertical grey line indicates the standard deviation, and the horizontal black dotted lines indicate the interquartile range.

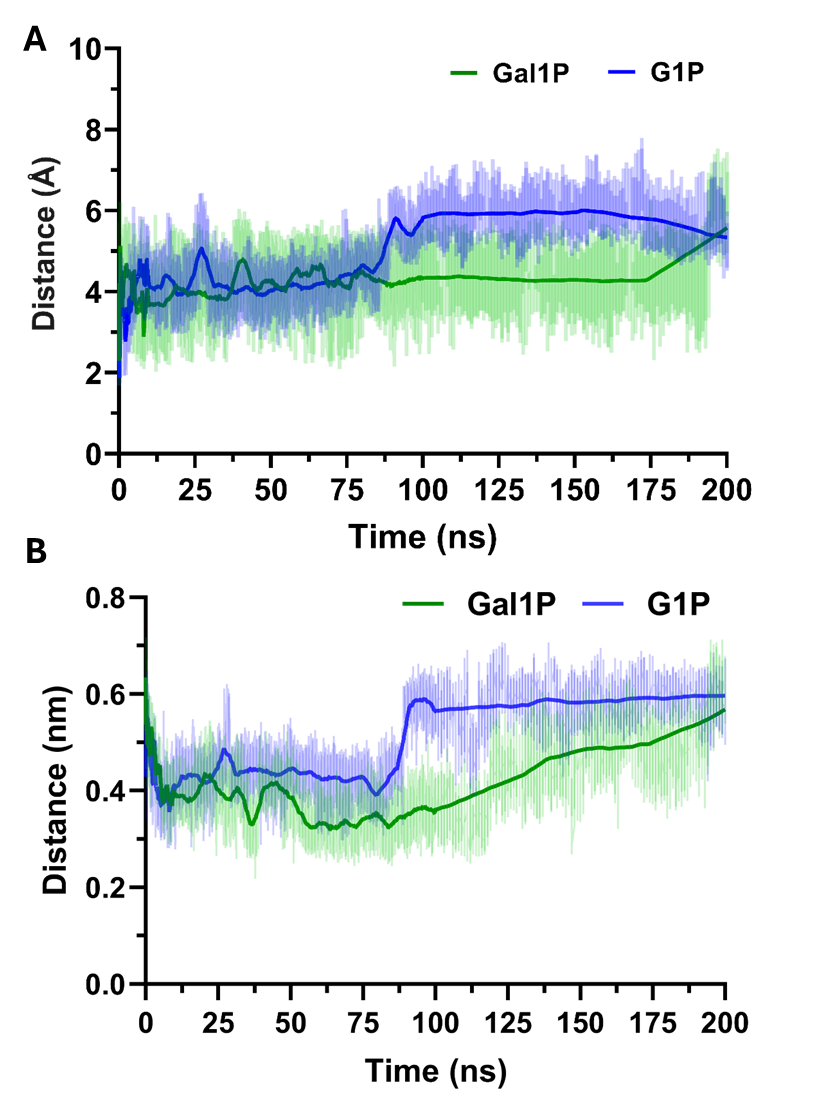

**Fig. S17.** Mean hydrogen bond distance between (**A**) E213 and 4OH of the substrates (Gal1P or G1P) and (**B**) N162 and 3OH of the substrates (Gal1P or G1P). The fits are plotted using 0^th^-order smoothing with 20 neighboring data points.

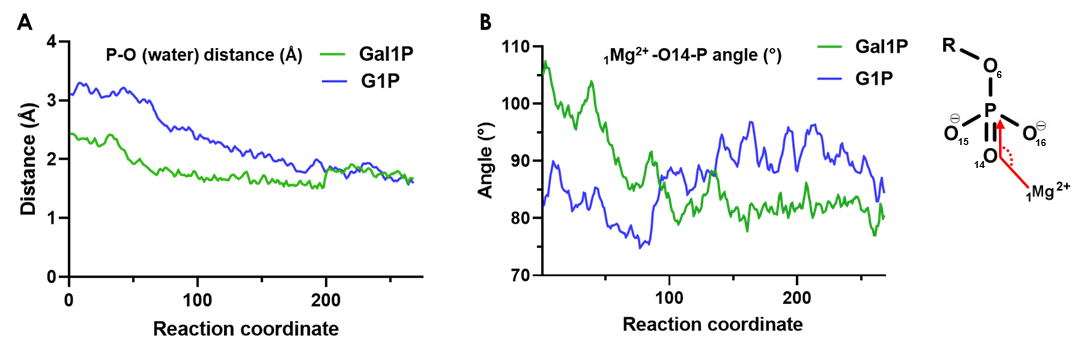

**Fig. S18.** QM/MM simulation data were used to evaluate (**A**) nucleophilic activation efficiency via phosphate–water distance and (**B**) catalytic geometry by the _1_Mg²⁺–O14–P angle. Here, O14 is one of the phosphoryl oxygen atom (i.e., the double-bonded oxygen on the phosphorus). This atom is located closest to _1_Mg²⁺ and the angle gives the geometrical alignment for the nucleophilic attack by hydroxyl ion to phosphate.

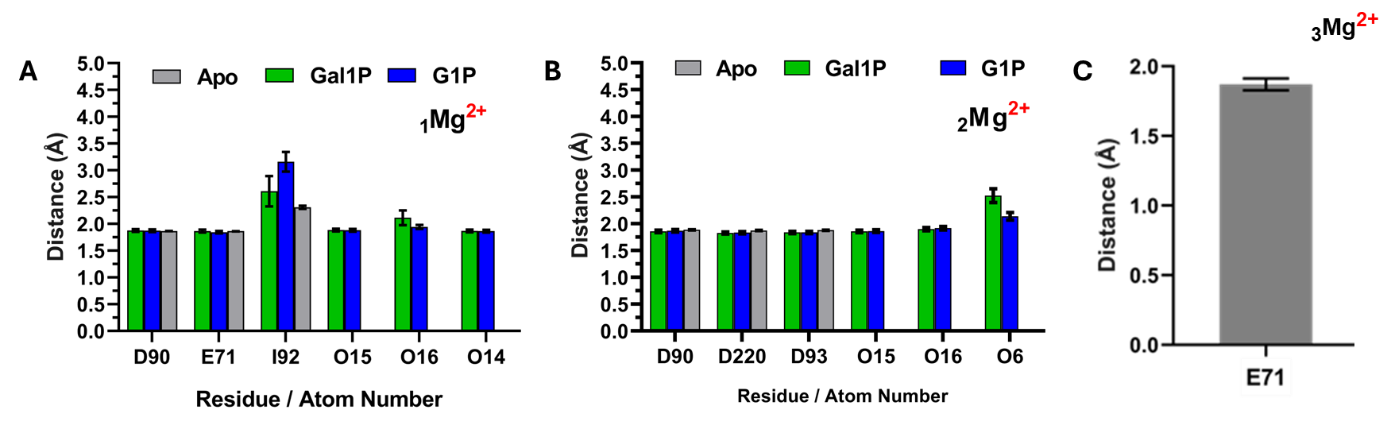

**Fig. S19:** The mean catalytic binding distances of Mg ions with the active site residues and substrates phosphate group atoms (**A**) _1_Mg^2+^ ion (**B**) _2_Mg^2+^ ion (**C**) _3_Mg^2+^ ion

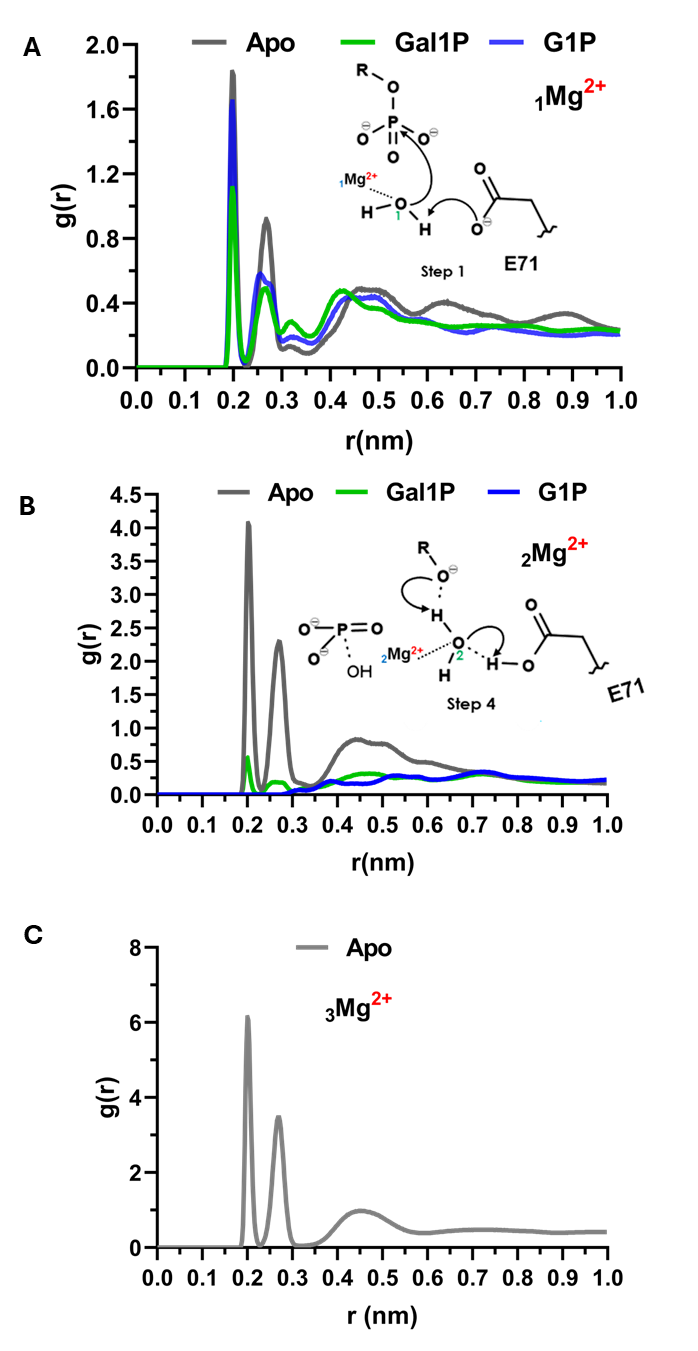
**Fig. S20:** Calculated radial distribution function (RDF) for _1_Mg^2+^, _2_Mg^2+^ and _3_Mg^2+^ ions to measure the probability density of finding a catalytic water molecule at a distance of **r** away from a selected Mg ions (**A**) _1_Mg^2+^ ion (**B**) _2_Mg^2+^ ion (**C**) _3_Mg^2+^ ion

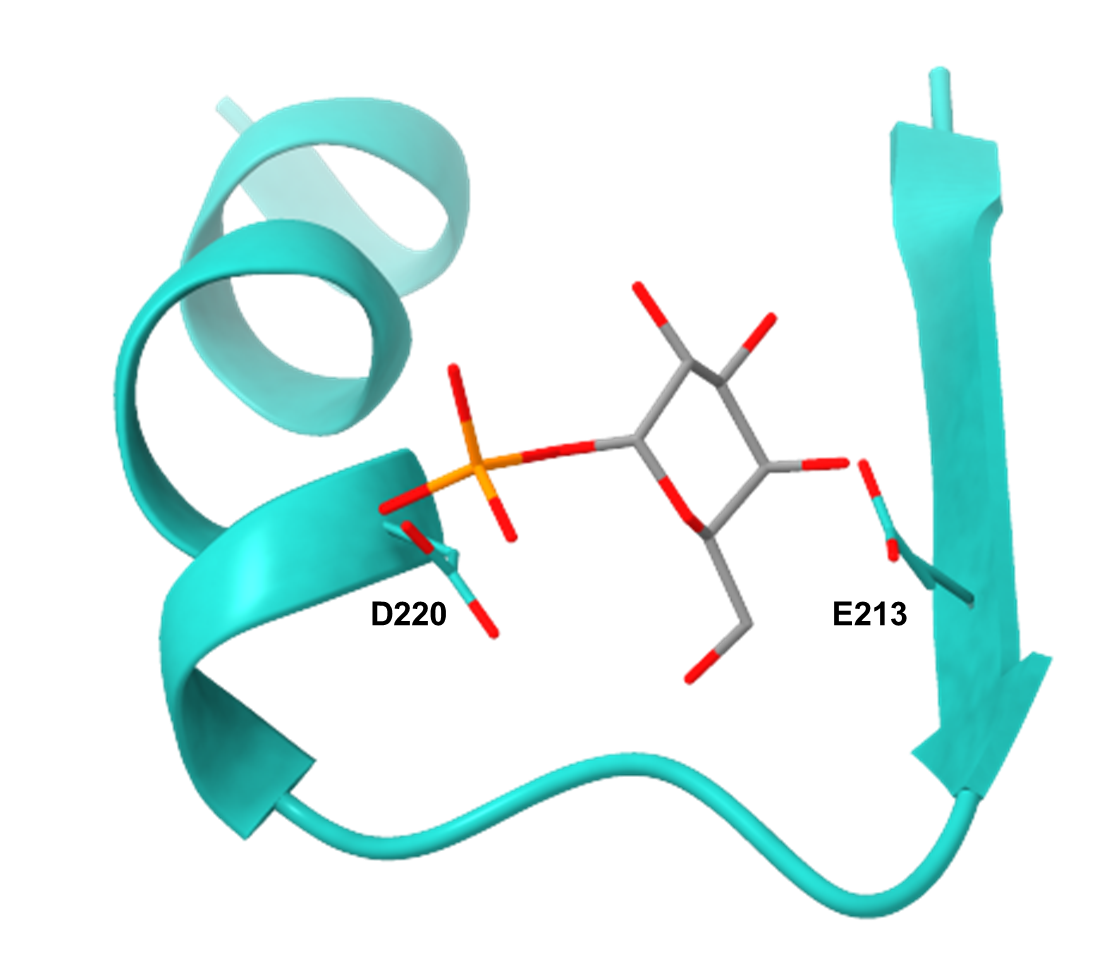

**Fig. S21:** Structural representation of the catalytic stabilization motif in DdGal1Pase. E213 is located on a β-sheet connected via a loop to an α-helix, where D220 is positioned. The α-helix spans residues 218–230, the β-sheet spans 210–214, and the loop region has a residue range of 215–217, respectively. Mg^2+^ ions were hidden for clarity.

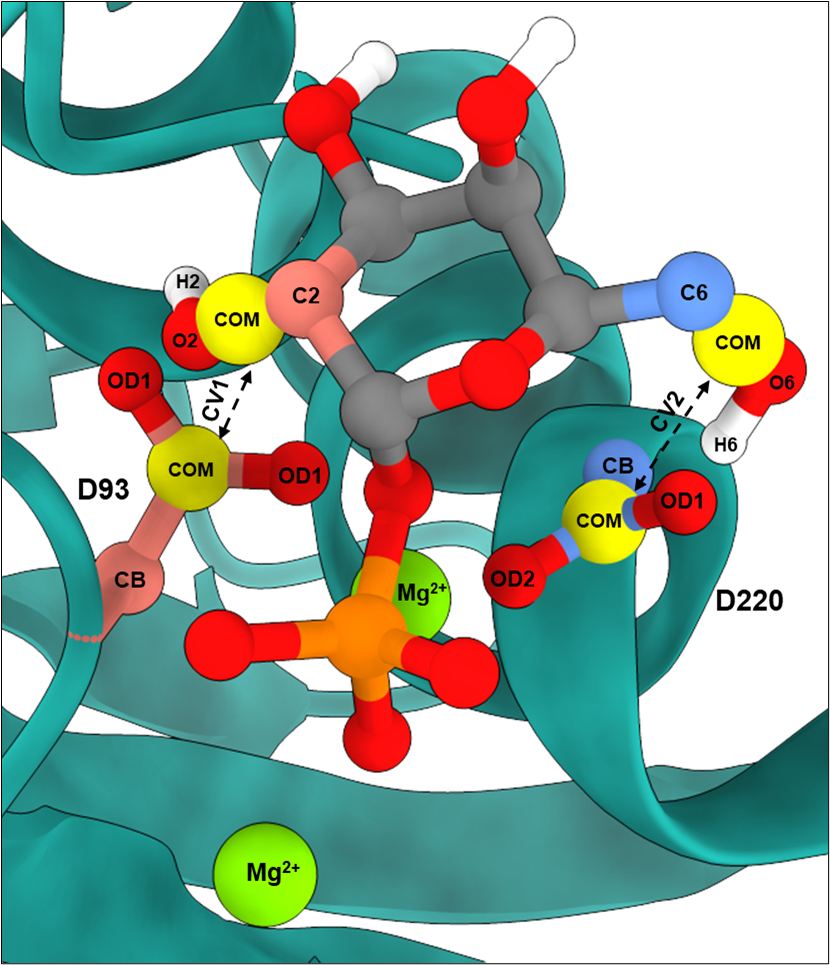

**Fig. S22.** Definition of collective variables (CVs) used in Metadynamics simulation. CV1 is defined as the distance between the center of mass (COM) of the side chain of D93, represented by the atoms Cβ (CB), Cγ (CG), OD1, and OD2, and the C2 hydroxyl (C2OH) group of Gal1P/G1P (Gal1P is shown for clarity). CV2 is defined as the distance between the COM of the side chain of D220, represented by the atoms Cβ (CB), Cγ (CG), OD1, and OD2, and the C6 hydroxyl (C6OH) group of Gal1P/G1P.

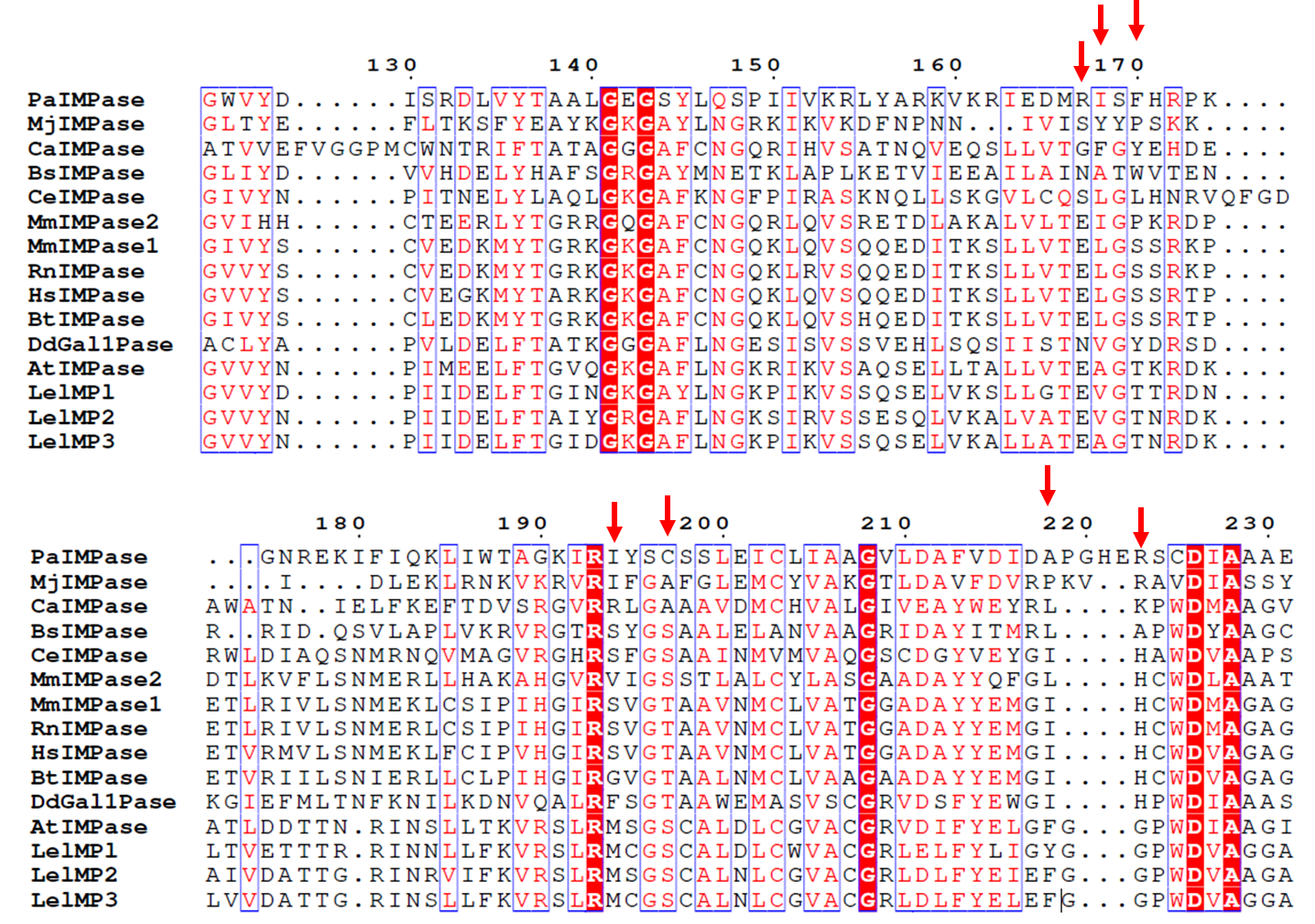

**Fig. S23.** The multiple sequence alignment of reported IMPases from *Lycopersicon esculentum (LeIMP1, Uniprot ID: P54926 , LeIMP2, Uniprot ID: P54927, LeIMP3, Uniprot ID: P54928)*, Homo sapiens (HsIMPase, PDB ID: 4AS4), *Arabidopsis thaliana* (AtIMPase, Uniprot ID: Q9M8S8), *Cicer arietinum* (CaIMPase, Uniprot ID:A0A1S2XI20), *Bos taurus* (BtIMPase, PDB ID: 2BJI), *Dictyostelium discoideum* (DdGal1Pase,Uniprot ID: Q54U72|), *Caenorhabditis elegans* (CeIMPase, Uniprot ID: Q19420), *Methanocaldococcus jannaschii* (MjIMPase, PDB ID: 1G0H), *Mus musculus* (MmIMPase1, Uniprot ID:AAB97469, MmIMPase2, Uniprot ID: AAK39516), *Rattus norvegicus* (RnIMPase, Uniprot ID:P97697), *Bacillus subtilis* (BsIMPase, Uniprot ID:CAB13340), *Pseudomonas arsenicoxydans* (PaIMPase, Uniprot ID: A0A4P6G7J6) having variations in the active site residues as compared to the DdGal1Pase marked with red arrows.

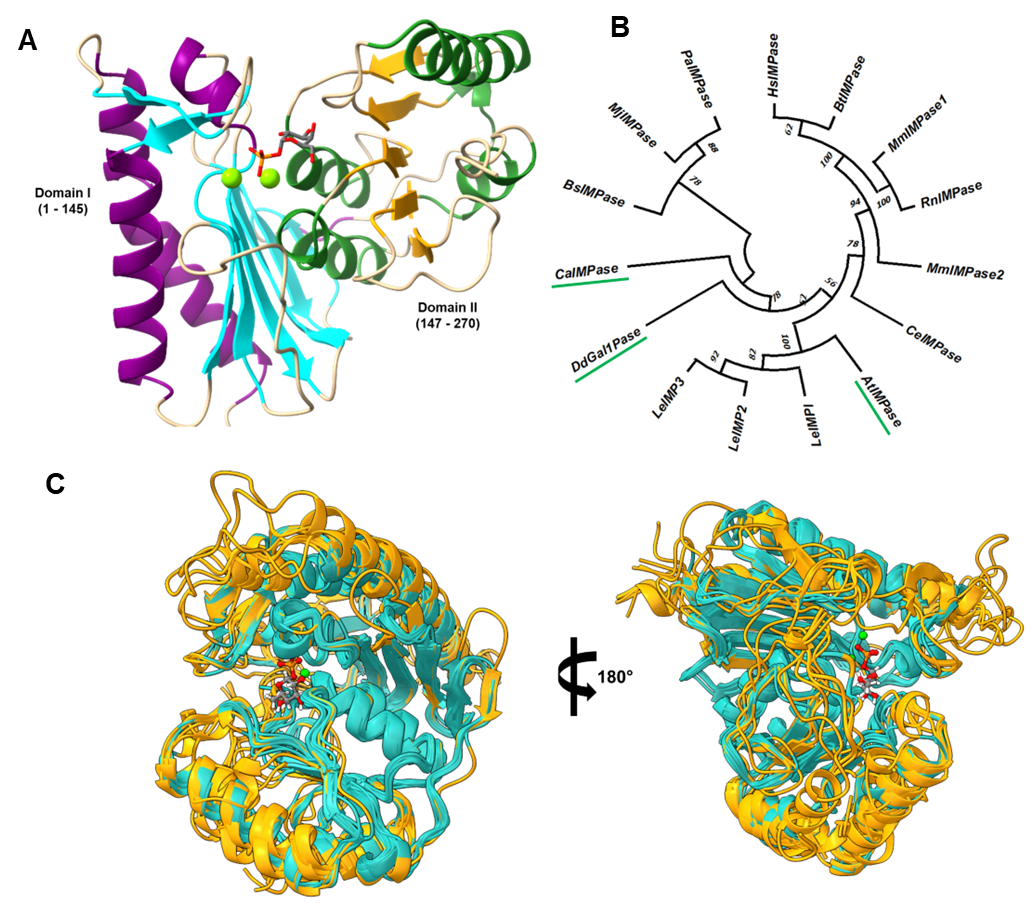

**Fig. S24. (A**) Cartoon diagram of the overall folding of the DdGal1Pase with indication of both the domains I and II. This depicts a penta-layered αβαβα sandwich. The α-helices belonging to domain I are indicated in purple while those belonging to domain II are indicated in forest green, respectively. Furthermore, the β-sheets set belonging to Domain I are indicated in cyan while those belonging to domain II are indicated in orange. Mg^2+^ ions are shown as green spheres. Representative Gal1P substrate is indicated in grey sticks. Domain analysis was conducted using InterProScan (Classification of Protein Families) (Matthias Blum, Antonina Andreeva , *et.al*, *Nucleic Acids Research* (2024), gkae1082, PMID: [39565202](https://europepmc.org/article/MED/39565202)) . (**B**) The evolutionary analysis of reported IMPases *Lycopersicon esculentum* (LeIMP1, Uniprot ID: P54926 , LeIMP2, Uniprot ID: P54927, LeIMP3, Uniprot ID: P54928), *Homo sapiens* (HsIMPase, PDB ID: 4AS4), *Arabidopsis thaliana* (AtIMPase, Uniprot ID: Q9M8S8), *Cicer arietinum* (CaIMPase, Uniprot ID:A0A1S2XI20), *Bos taurus* (BtIMPase, PDB ID: 2BJI), *Dictyostelium discoideum* (DdGal1Pase,Uniprot ID: Q54U72|), *Caenorhabditis elegans* (CeIMPase, Uniprot ID: Q19420), *Methanocaldococcus jannaschii* (MjIMPase, PDB ID: 1G0H), *Mus musculus* (MmIMPase1, Uniprot ID:AAB97469, MmIMPase2, Uniprot ID: AAK39516), *Rattus norvegicus* (RnIMPase, Uniprot ID:P97697), *Bacillus subtilis* (BsIMPase, Uniprot ID:CAB13340), *Pseudomonas arsenicoxydans* (PaIMPase, Uniprot ID: A0A4P6G7J6) having activity for inositol, while the green underlined IMPases have shown activity for Gal1P; The evolutionary history was inferred by using the Maximum Likelihood method and Whelan and Goldman model. A discrete Gamma distribution was used to model evolutionary rate differences among sites (5 categories (+G, parameter = 2.2637)) (**C**) The structural superimposition of all above IMPases over DdGal1Pase, where the RMSD below 2 are colored in light sea green color, while the RMSD above 2 are colored in the orange color.

**Table S1.** Transport-associated genes targeted for deletion to study selective partitioning of galactose and tagatose across the cell membrane to improve equilibrium conversion.

| **#** | **Gene** | **Function** | **Reference** |
| --- | --- | --- | --- |
| **1** | *setA* | Sugar exporter | Carreón-Rodríguez et al. 2023 |
| **2** | *setB* | Sugar exporter | Carreón-Rodríguez et al. 2023 |
| **3** | *setC* | Arabinose exporter | Koita et al. 2012 |
| **4** | *mglBAC* | Galactose importer | Carreón-Rodríguez et al. 2023, Kim et al. 2008 |
| **5** | *galP* | Galactose importer | Carreón-Rodríguez et al. 2023 |
| **6** | *lacY* | Lactose importer | Tong et al. 2020 |
| **7** | *araE* | Arabinose importer | Carreón-Rodríguez et al. 2023 |
| **8** | *ydeA* | Implicated in arabinose efflux | Koita et al. 2012 |
| **9** | *ydhC* | Implicated in arabinose efflux | Koita et al. 2012 |
| **10** | *nagC* | Represses galactose transport | El Qaidi et al. 2009 |
| **11** | *galR* | Galactose catabolism repressor | Carreón-Rodríguez et al. 2023 |
| **12** | *galS* | Galactose catabolism repressor | Carreón-Rodríguez et al. 2023 |
| **13** | *ydeE* | Implicated in arabinose efflux | Carreón-Rodríguez et al. 2023 |

**Table S2.** Mean hydrogen bond occupancy and standard error of mean of Gal1P and G1P with active site residues. Cryptic Hydrogen bond occupancy refers to the mean occupancy of hydrogen bonds that have less than 1% occupancy across the 3 × 200 ns simulations.

|  | **Gal1P** | | **G1P** | |
| --- | --- | --- | --- | --- |
| **Residue** | **Mean occupancy (%)** | **Standard error** | **Mean occupancy (%)** | **Standard error** |
| D220 | 42.983 | 6.902 | 22.817 | 16.900 |
| N162 | 16.237 | 11.126 | 11.333 | 5.866 |
| G94 | 8.897 | 0.447 | 13.963 | 1.487 |
| G194 | 8.293 | 3.388 | 3.963 | 2.301 |
| T95 | 6.353 | 5.537 | 6.163 | 6.158 |
| E213 | 3.620 | 3.152 | 1.327 | 1.019 |
| K38 | 2.767 | 1.869 | 2.927 | 1.456 |
| D93 | 2.323 | 1.070 | 22.200 | 7.256 |
| G215 | 1.290 | 0.283 | 2.587 | 1.797 |
| Y165 | 0.887 | 0.782 | 0.067 | 0.037 |
| A196 | 0.137 | 0.050 | 0.720 | 0.508 |
| W219 | 0.053 | 0.028 | 0.333 | 0.313 |
| D42 | 0.023 | 0.012 | 0.587 | 0.358 |
| K75 | 0.003 | 0.003 | 0.000 | 0.000 |
| H217 | 0.000 | 0.000 | 0.037 | 0.020 |
| T96 | 0.000 | 0.000 | 0.010 | 0.010 |
| G164 | 0.000 | 0.000 | 0.003 | 0.003 |

**Table S3.** Number of atoms in molecular simulation for different systems in this study. Each complex structure was simulated three times for 200 ns, totaling 1.2 µs of simulation time.

| **Molecule type** | **DdGal1Pase-Gal1P** | **DdGal1Pase-G1P** | **DdGal1Pase (apo)** |
| --- | --- | --- | --- |
| Protein | 4,200 | 4,200 | 4,200 |
| Water | 18,807 | 20,202 | 18,831 |
| Na^+^ | 29 | 32 | 25 |
| Cl^-^ | 22 | 25 | 22 |
| Mg^2+^ | 2 | 2 | 3 |
| Substrate | 27 | 27 | NA |
| Total Atoms | 23,087 | 24,488 | 23,081 |
